# Estimating the contribution of musculoskeletal impairments to altered gait kinematics in children with cerebral palsy using predictive simulations

**DOI:** 10.1101/2025.06.30.661463

**Authors:** Bram Van Den Bosch, Lars D’Hondt, Ilse Jonkers, Kaat Desloovere, Anja Van Campenhout, Friedl De Groote

## Abstract

1.

**Background:** Cerebral palsy (CP) is caused by a brain lesion around birth leading to impaired motor control, bony deformities, muscle contractures, and weakness resulting in altered gait. Since the brain lesion cannot be cured, treatment aims at improving mobility. Multilevel surgery targets muscle and bony impairments but the outcome of multilevel surgery is variable and unpredictable due to our limited insight in the contribution of musculoskeletal impairments to gait alterations.

**Methods:** Here, we used predictive simulations based on personalized musculoskeletal models to identify the contribution of musculoskeletal impairments to altered gait in eight individuals with CP scheduled for multilevel surgery. For each individual, we generated gait patterns based on eight models with different levels of personalization. We modeled muscle weakness, muscle contractures, and/or bony deformities of hip and knee and evaluated the contribution of these impairments to deviations in kinematics by comparing simulated and experimental kinematics.

**Results:** The contribution of musculoskeletal impairments varied largely between participants. All modeled musculoskeletal impairments explained little to 39% of the kinematic deficits, in line with the limited and variable effect of multilevel surgery. Muscle contractures had the largest effect on the predicted kinematics.

**Conclusion:** Our results suggest an important contribution of motor control and unmodeled musculoskeletal impairments (e.g. shank and foot deformities) to alterations in the gait pattern. Model-based simulations are a promising tool to determine the contribution of musculoskeletal impairments to alterations in gait kinematics in individuals with CP.

## 2. Background

Cerebral palsy (CP) is one of the most common neurological disorders in children with an estimated prevalence of 2‰ – 2.3‰ live births [1]. CP is caused by a non-progressive brain lesion around birth, which causes motor control impairments and due to altered muscle control and skeletal loading also musculoskeletal (MSK) impairments [2]. Common musculoskeletal impairments are bony deformities and muscle contractures [3]. As the brain lesion cannot be cured, treatment aims at improving mobility. Multilevel surgery (MLS) is common in CP and targets multiple MSK impairments [4]. Muscle length is corrected by muscle-tendon lengthening procedures and bony deformities are corrected using derotation osteotomies. Unfortunately, the effect of MLS on walking kinematics is variable and improvements in kinematics (main outcome) have stagnated over the last two decades [5]. Predicting the outcome of MLS, and hence avoiding unsuccessful surgeries, is challenging due to the complex interactions between the many different motor control and musculoskeletal impairments. Here, we aimed at isolating the contribution of different musculoskeletal impairments, targeted by MLS, to alterations in walking kinematics in CP.

Model-based simulations of walking are a powerful tool to study the contribution of MSK impairments to gait deficits [6]. Such simulations generate novel walking patterns based on a musculoskeletal model without relying on experimental gait data. It is typically assumed that humans walk in a way that minimizes a movement-related cost. As a result, muscle excitations and the resulting walking pattern can be found by solving an optimal control problem. Such simulations yield muscle excitations and joint kinematics of healthy walking that are in close agreement with experimental observations [7], [8]. As these simulations do not rely on experimental data to generate novel walking patterns, they can be used to study the effect of alterations in musculoskeletal properties on the walking pattern by changing model parameters and re-solving the optimal control problem.

Predictive simulations have been used to study the isolated effect of common musculoskeletal impairments on walking kinematics in CP. Ong et al. [9] and Bruel et al. [10] used simulations to isolate the effect of ankle plantar flexor weakness, contractures, and hyper-reflexia - common impairments in CP – on gait kinematics. Plantar flexor weakness in isolation led to heel-walking, whereas plantar flexor contractures and hyper-reflexia led to toe-walking. Yet, so far, only one case study has used simulations to isolate the contributions of musculoskeletal impairments in a specific child with CP. Falisse et al. performed simulations based on a personalized neuro-musculoskeletal model to dissociate the contributions of musculoskeletal and motor control impairments to deviations in gait kinematics in a child with CP scheduled for MLS [11]. In this child, altered musculoskeletal geometry and muscle-tendon properties rather than motor control deficits were the primary cause of the observed crouch gait pattern. Given the heterogeneity of musculoskeletal impairments in children with CP, results from a single case study cannot be generalized. Therefore, we sought to characterize the contribution of musculoskeletal impairments across a group of individuals with CP scheduled for MLS.

Several methods for model personalization have been developed. Scheys et al. developed a method to derive musculoskeletal geometry from magnetic resonance images (MRI) [12]. They showed that personalizing joint locations and muscle-tendon paths based on MRI considerably affected the moment arms in children with increased femoral anteversion. Personalized modeling resulted in smaller moment arms for hip flexors, extensors, abductors, adductors and external rotators but larger moment arms for hip internal rotators [13]. Moment arms co-determine the torque generating capacity of the muscles and might thereby affect the predicted gait pattern. Muscle properties can be further personalized by fitting the measured and simulated torque-angle relationship across a range of movements [14], [15], [16], [17], [18], [19]. However, accurate parameter estimation requires sufficiently rich experimental data (spanning a range of motions) [19], whereas typically only gait is assessed during a clinical gait analysis. On the other hand, an extensive clinical examination is often performed as part of treatment planning, including tests to evaluate muscle length, strength, alignment and motor control. Whereas the clinical examination has an established clinical relevance, data collected during the examination has not been used to personalize models.

The aim of this study was to determine to what extent musculoskeletal impairments contribute to altered gait in children with CP who are scheduled for MLS. To this aim, we performed predictive simulations of walking for eight individuals with CP. We used eight models per individual with different levels of personalization to evaluate the contribution of muscle weakness, muscle contractures and bony deformities. Muscle weakness and contractures were modeled based on the clinical examination and bony deformities were modeled based on MRI. We evaluated the agreement between experimental and simulated kinematics for each individual and model. We determined the contribution of the modeled impairments by comparing the difference between experimental and simulated kinematics for simulations based on a scaled generic model and the personalized models.

## 3. Methods

### 3.1 Description of study participants

Data were collected for eight individuals with spastic CP planned for MLS at the time of their pre-operative clinical gait assessment (Table 1). Parents or caregivers signed an informed consent form, participants older than twelve signed an informed assent form. The study was approved by the Ethical Committee UZ/KU Leuven (S64909).

**Table 1.** Study participants.

| ID | Sex | Topography<sup>a</sup> | Age<br>(years) | GMFCS<sup>b</sup> | Body height<br>(cm) | Body mass<br>(kg) | Walking speed<sup>c</sup><br>(m/s) |
| --- | --- | --- | --- | --- | --- | --- | --- |
| CP1 | M | Bi | 12 - 13 | 2 | 145 - 150 | 30 - 35 | 1.33 |
| CP2 | M | Bi R | 16 - 17 | 1 | 170 – 175 | 55 - 60 | 1.07 |
| CP3 | F | Uni R | 15 - 16 | 1 | 165 - 170 | 55 - 60 | 1.17 |
| CP4 | M | Bi L | 17 - 18 | 2 | 170 – 175 | 45 - 50 | 0.61 |
| CP5 | M | Asym L | 16 - 17 | 2 | 170 - 175 | 50 - 55 | 1.05 |
| CP6 | M | Bi | 15 - 16 | 2 | 160 – 165 | 40 - 45 | 0.99 |
| CP7 | M | Uni L | 21 - 22 | 1 | 165 - 170 | 45 - 50 | 1.02 |
| CP8 | F | Bi | 10 - 11 | 2 | 130 – 135 | 25 - 30 | 0.83 |

*Ranges for age, body height and body mass were reported to improve anonymization.*
<sup>a</sup> *Bi = bilateral CP, Bi R = bilateral CP with right side most affected, Bi L = bilateral CP with left side most affected, Uni R = Unilateral CP right, Uni L = Unilateral CP left, Asym L = Asymmetric bilateral CP with left arm.*
<sup>b</sup> *Gross Motor Function Classification System*
<sup>c</sup> *Average self-selected walking speed*

### 3.2 Description of experimental data

All study participants had a clinical examination and 3D gait analysis followed by an MRI scan of the legs and pelvis.

#### Clinical examination

The clinical examination was performed by an experienced clinician. This examination was part of standard clinical care and was used to inform clinical decision making. It assessed muscle length, strength, bony alignment, and motor control. Muscle length was evaluated by passively moving the joints to the end of range of motion and measuring the final angle with a goniometer. Evaluated joint motions were hip flexion, hip extension, knee flexion, knee extension, ankle dorsiflexion, and ankle plantarflexion. Muscle strength was evaluated using modified Manual Muscle Testing (MMT). The strength of hip flexors, extensors, abductors and adductors, knee flexors and extensors, ankle plantar flexors and dorsiflexors, invertors and evertors was tested. When no muscle contraction could be palpated, the muscle group was scored 0, when the individual could move against gravity with maximal resistance the muscle group was scored 5 (see supplement, Table S2). A specific evaluation of abdominal and back muscles was also performed and scored between 0 and 5 (see supplement, Table S3 and S4). For alignment, femoral anteversion [20], bimalleolar angle and tibiofemoral angle [21] were evaluated. For motor control, both spasticity and selectivity were assessed. Spasticity was assessed using the modified Ashworth scale [22], Tardieu scale [23] and Duncan-Ely [21]. The Selective Motor Control Test [24] and Selective Control Assessment of the Lower Extremity [25] were used to evaluate selectivity.

#### 3D Gait Analysis

The gait analysis was performed at a self-selected speed and marker trajectories were captured with a 12-camera Vicon system (Vicon, Oxford, UK; sampling frequency of 100 Hz). Markers were attached according to an extended lower limb plug-in gait model (see supplement, Table S1). Ground reaction forces were collected at 1000 Hz with two embedded force plates (AMTI, Watertown MA, USA) in the raised 10-meter walkway. Muscle activity of eight muscles was measured bilaterally with a 16-channel telemetric surface electromyography system (Zerowire, Cometa, Italy) at 1000 Hz. The electrodes were placed according to the SENIAM guidelines [26]. In this study, we only evaluated and compared kinematics.

#### MRI

MRI of the lower limbs and pelvis were acquired similar to Bosmans et al. [27]. Depending on the individual’s size, three to five axial image series were acquired on a 3T Siemens MR scanner using a T1 weighted SE sequence with the participants lying supine with extended knees. For the image series containing the hip, knee, or ankle, inter-slice distance was 1 mm with a voxel size of 1.04 x 1.04 x 1 mm. For the other image series, the inter-slice distance was 2 mm with a voxel size of 1.04 x 1.04 x 2 mm. Glycerin markers were placed on the marker locations of the 3DGA to ensure correct registration for inverse kinematics.

### 3.3 Predictive simulations

We performed predictive simulations of walking using PredSim [7], [28]. In short, we solved for the gait cycle duration, muscle controls, and the corresponding gait pattern by minimizing a cost function while imposing task constraints and musculoskeletal dynamics without relying on experimental gait data. The task constraints were an imposed average forward speed of the pelvis as well as periodicity. We prescribed the participants’ walking speed in the simulations. Since we did not explicitly model interactions between limbs, we used distance constraints scaled based on body height to prevent segments penetrating each other. We used a previously determined cost function [7], i.e. the integral of the weighted sum of squared metabolic energy rate (*Ė*), muscle activations (*a*), joint accelerations (*u*_*a*_), and passive joint torques (*T*_*p*_):

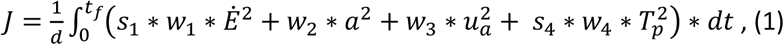

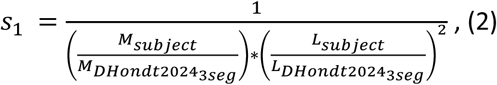

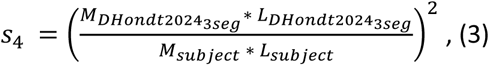

where *d* is the distance traveled, *t*_*f*_ is gait cycle duration, *t* is time, and *w*_1_ − *w*_4_ are weight factors, *M*_*subject*_ is the mass of the subject, *L*_*subject*_ is the height of the subject, and 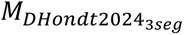 and 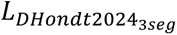 are the mass and height of the generic model (described in detail below). Muscle metabolic energy was calculated using the model of Bhargava et al. [29], which was made continuously differentiable by approximating conditional statements with a hyperbolic tangent.

The resulting optimal control problems were solved using direct collocation (100 mesh intervals, 3 collocation points) and algorithmic differentiation. Problems were formulated and solved in MATLAB. Skeleton dynamics was formulated through OpenSimAD [30], CasADi [31] was used for problem formulation and algorithmic differentiation, and the resulting nonlinear programming problems were solved in IPOPT [32] (tolerance: 10^-4). For each model, we used at least two initial guesses. The converged simulation with the lowest cost was chosen as final result.

### 3.4 Model personalization

We performed a series of simulations based on models with different levels of personalization to evaluate how musculoskeletal impairments affect the gait pattern. We divided the musculoskeletal impairments into muscle weakness, muscle contractures, and bony deformities. Muscle weakness and contractures were derived from the clinical examination. We derived bony deformities from the MR images using a previously developed approach to determine hip joint centers and knee axes as well as muscle-tendon paths of muscles spanning the hip and knee (but not the ankle) [12]. To investigate the contribution of these impairments as well as interaction effects, we created eight musculoskeletal models as described in Table 2.

**Table 2.** Model description.

|  | Model |  |  |  |  |  |  |  |
| --- | --- | --- | --- | --- | --- | --- | --- | --- |
|  | GEN | GENWEAK | GENCTR | GENFULL | GEO | GEOWEAK | GEOCTR | GEOFULL |
| **Modeled impairment** |  |  |  |  |  |  |  |  |
| *Muscle weakness<sup>a</sup>* |  | x |  | x |  | x |  | x |
| *Muscle contractures<sup>b</sup>* |  |  | x | x |  |  | x | x |
| *Bony deformities<sup>c</sup>* |  |  |  |  | x | x | x | x |

<sup>a</sup> Muscle weakness was modeled as reduced active fiber force with scaling factors determined based on Manual Muscle Testing scores. When combined with bony deformities, the scaling factor was adjusted to account for the contribution of moment arm deficits to weakness.
<sup>b</sup> Muscle contractures were modeled by reducing optimal fiber length to represent the reduced number of sarcomeres in series that has been observed in CP. Coordinate limit torques for knee extension were adjusted to reflect the knee extension deficit. Coordinate limit torques for plantar flexion were adjusted to reflect plantar flexion deficit if present.
<sup>c</sup> Bony deformities of hip and knee as well as alterations in the paths of muscles spanning the hip and knee were included using an established workflow for MRI-based modeling.

#### Generic musculoskeletal model

All models were based on the model proposed by D’Hondt et al. [33]. The model has 31 degrees of freedom (DOFs) (pelvis-to-ground: 6 DOFs, hip: 3 DOFs, knee: 1 DOF, ankle: 1 DOF, subtalar: 1 DOF, metatarsophalangeal: 1 DOF, lumbar: 3 DOFs, shoulder: 3 DOFs, and elbow: 1 DOF). The lower limb and lumbar joints are actuated by 92 Hill-type muscle-tendon units [34], [35]. The shoulder and elbow joints are actuated by eight ideal torque actuators. Foot-ground contact is modeled by five Hunt-Crossley contact spheres per foot. Passive joint torques with exponential stiffness and damping [36] were added to the lower limb and lumbar joints to represent the effects of unmodeled passive structures [7]. Muscle excitation-activation coupling was described by Raasch’s model [37], [38]. Skeletal motion was modeled with Newtonian rigid body dynamics [30], [39]. We removed quadratus femoris and gemellus because modeling bony deformities caused unrealistic operating ranges (>1.5 normalized fiber lengths) of these small muscles for some participants whereas removing both muscles from the generic model had little effect on the simulated gait pattern.

#### Reference model

The reference model (GEN) is the model proposed by D’Hondt et al. [33] scaled to the participant’s anthropometry using the OpenSim Scale Tool [40]. We additionally scaled maximal isometric force, which is not affected by scaling in OpenSim, based on subject mass [41]:

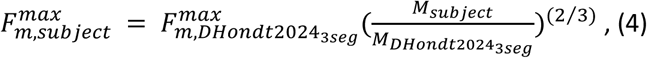

With *F*^*max*^ the maximal isometric force and *M* the mass of the respective models. Parameters of the foot-ground contact model were also scaled based on the subject’s anthropometry (see supplement, equations s1-s3). This model does not include impairments, with exception of leg length differences and was used as the basis for further personalization.

#### Modeling muscle weakness

To model muscle weakness, active fiber force was scaled based on the MMT scores from the clinical examination:

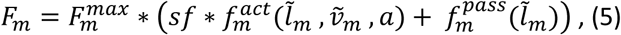

where *F*_*m*_ is muscle force, 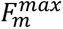 is the maximal isometric force, *sf* is the weakness scale factor, 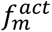 is the active muscle force-length-velocity characteristic, 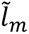 is normalized fiber length, 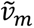 is normalized fiber velocity, *a* is muscle activation, and 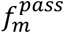 is the passive muscle force-length characteristic. The weakness scale factors were determined semi-arbitrarily. We performed preliminary simulations with different sets of scaling factors (i.e. [1.0, 0.7, 0.5, 0.3, 0.2, 0.1] and [0.7, 0.5, 0.3, 0.2, 0.1, 0.05] for MMT scores of 5, 4, 3, 2, 1, 0 combined with either the same of larger reductions in strength for plantar flexors) in a subset of the participants (four out of eight) and selected the scale factors for which weakness explained most of the participants’ gait deficits. This resulted in scale factors for active fiber force of 0.7, 0.5, 0.3, 0.2, 0.1, and 0.05 for MMT scores of 5, 4, 3, 2, 1, 0, respectively. For plantar flexors, we reduced the scaling factor by one step (e.g. a score of 3 corresponds to a scaling factor of 0.2 instead of 0.3).

We grouped muscles to link the MMT scores to individual muscles. Hip muscles were divided into abductors (gluteus minimus and medius, tensor fascia latea and piriformis), adductors (adductors and pectineus), flexors (iliacus and psoas) and extensors (gluteus maximus and biceps femoris long head). Knee muscles were divided into flexors (remaining knee flexors attaching on pelvis or femur) and extensors (quadriceps). Muscles spanning the ankle were divided into plantar flexors (gastrocnemius medialis, gastrocnemius lateralis, soleus), dorsiflexors (tibialis anterior and toe extensor muscles), evertors (peronei) and invertors (toe flexors and tibialis posterior).

#### Modeling muscle contractures

Muscle fiber lengths have been observed to be shorter in CP [42]. Therefore, we chose to model contractures in the Hill-type muscles by reducing optimal fiber length. When optimal fiber length is reduced, muscle fibers will be stretched more at the same muscle-tendon length resulting in higher passive forces.

We only modeled contractures when there was a clinical indication (Table 3), i.e. clinical examination angles for passive range of motion were out of the typical range. To determine the optimal fiber length for contracted muscles, we placed the musculoskeletal model in the same position as during the passive range of motion assessment and activated the muscle to 1%. We then determined the optimal fiber length such that the net modeled joint torque was 15 Nm at the end of range of motion. The threshold of 15 Nm was selected based on data from an instrumented measurement using a handheld force cell of one of the participants. We performed a sensitivity analysis on a subset of the participants where the threshold was set to 10 Nm. Simulations with models based on the lower threshold explained less of the kinematic deficits than simulations with models based on the 15 Nm threshold.

**Table 3.** Clinical examination.

|  | CP1 |  | CP2 |  | CP3 |  | CP4 |  | CP5 |  | CP6 |  | CP7 |  | CP8 |  |
| --- | --- | --- | --- | --- | --- | --- | --- | --- | --- | --- | --- | --- | --- | --- | --- | --- |
|  | L | R | L | R | L | R | L | R | L | R | L | R | L | R | L | R |
| **Passive Range of Motion** |  |  |  |  |  |  |  |  |  |  |  |  |  |  |  |  |
| Hip |  |  |  |  |  |  |  |  |  |  |  |  |  |  |  |  |
| *Flexion* | 0 | 0 | 0 | 0 | 0 | 0 | 0 | 0 | 0 | 0 | 0 | 0 | 0 | 0 | 0 | 0 |
| *Extension* | 0 | 0 | 0 | 0 | 0 | 0 | -10<sup>a</sup> | 0 | 0 | 0 | 0 | 0 | 0 | 0 | 0 | 0 |
| Knee |  |  |  |  |  |  |  |  |  |  |  |  |  |  |  |  |
| *Extension\** | 0<sup>a</sup> | 5<sup>a</sup> | 0<sup>a</sup> | -10<sup>c</sup> | 0<sup>a</sup> | 0<sup>a</sup> | -5<sup>b</sup> | 0<sup>a</sup> | -10<sup>c</sup> | -5<sup>b</sup> | -10<sup>c</sup> | -10<sup>c</sup> | 0<sup>a</sup> | 0<sup>a</sup> | -5<sup>b</sup> | -5<sup>b</sup> |
| *Spontaneous* | 0 | 0 | 0 | -20<sup>c</sup> | 0 | 0 | -20<sup>c</sup> | 0 | -15<sup>b</sup> | -10<sup>b</sup> | -15<sup>b</sup> | -25<sup>c</sup> | -10<sup>b</sup> | 0 | -10<sup>b</sup> | -15<sup>b</sup> |
| *Rectus length\** | 2<sup>b</sup> | 2<sup>b</sup> | 0 | 1<sup>a</sup> | 0 | 1<sup>a</sup> | 0 | 2<sup>b</sup> | 0 | 0 | 0 | 0 | 1<sup>a</sup> | 1<sup>a</sup> | 0 | 0 |
| *Patella alta* | 0 | 0 | 10<sup>a</sup> | 15<sup>b</sup> | 10<sup>a</sup> | 10<sup>a</sup> | 10<sup>a</sup> | 0 | 5<sup>a</sup> | 5<sup>a</sup> | 0 | 0 | 10<sup>a</sup> | 10<sup>a</sup> | 0 | 0 |
| *Popliteal angle (uni)\** | -65 | -60 | -65 | -65 | -55 | -60 | -85<sup>a</sup> | -75<sup>c</sup> | -85 | -70 | -65 | -80<sup>b</sup> | -65<sup>a</sup> | -60<sup>a</sup> | -80<sup>b</sup> | -80<sup>a</sup> |
| *Popliteal angle (bi)\** | -65<sup>b</sup> | -60<sup>b</sup> | -65<sup>b</sup> | -65<sup>b</sup> | -55<sup>b</sup> | -60<sup>b</sup> | -80<sup>c</sup> | -60<sup>b</sup> | -85<sup>c</sup> | -70<sup>c</sup> | -65<sup>b</sup> | -70<sup>c</sup> | -60<sup>b</sup> | -55<sup>b</sup> | -70<sup>c</sup> | -75<sup>c</sup> |
| Ankle |  |  |  |  |  |  |  |  |  |  |  |  |  |  |  |  |
| *Dorsiflex (90°)\** | 20 | 25 | 15<sup>a</sup> | 0<sup>b</sup> | 10<sup>a</sup> | 0<sup>b</sup> | 5<sup>b</sup> | 20 | 25 | 20 | 30 | 35 | -5<sup>c</sup> | 10<sup>a</sup> | 20 | 15<sup>a</sup> |
| *Dorsiflex (0°)\** | 10 | 15 | 10 | -5<sup>b</sup> | 5<sup>a</sup> | 0<sup>a</sup> | 0<sup>a</sup> | 10 | 10 | 10 | 10 | 15 | -10<sup>c</sup> | 0<sup>a</sup> | 10 | 10 |
| **Alignment** |  |  |  |  |  |  |  |  |  |  |  |  |  |  |  |  |
| *Anteversion angle* | 30<sup>a</sup> | 35<sup>a</sup> | 15 | 25<sup>a</sup> | 10 | 15 | 15 | 15 | 25<sup>a</sup> | 10 | 25<sup>a</sup> | 20 | 35<sup>c</sup> | 25<sup>a</sup> | 30<sup>a</sup> | 30<sup>a</sup> |
| *Tibiofemoral angle* | 5<sup>a</sup> | 0<sup>a</sup> | 15 | 10 | 15 | 10 | 15 | 15 | 20 | 10 | 0<sup>a</sup> | 15 | -15<sup>c</sup> | 5<sup>a</sup> | 5<sup>a</sup> | 15 |
| *Bimalleolar angle* | 10 | 10 | 15 | 10 | 15 | 10 | 15 | 15 | 10 | 5<sup>a</sup> | 10 | 20 | 10 | 15 | 15 | 15 |
| **Motor control** |  |  |  |  |  |  |  |  |  |  |  |  |  |  |  |  |
| *Selective total (max = 22)* | 19,5<sup>a</sup> | 21,5<sup>a</sup> | 22 | 21,5<sup>a</sup> | 22 | 22 | 16,5<sup>b</sup> | 21<sup>a</sup> | 13,5<sup>c</sup> | 22 | 19<sup>a</sup> | 16<sup>b</sup> | 13,5<sup>c</sup> | 19<sup>a</sup> | 16,5<sup>b</sup> | 14<sup>c</sup> |
| *SCALE total (max = 10)* | 8<sup>a</sup> | 9<sup>a</sup> | 10 | 10 | 10 | 10 | 6<sup>b</sup> | 10 | 6<sup>b</sup> | 9<sup>a</sup> | 7<sup>b</sup> | 6<sup>b</sup> | / | / | 9<sup>a</sup> | 7<sup>b</sup> |
| *Spasticity\*\* (max = 32)* | 5<sup>b</sup> | 3<sup>b</sup> | 3<sup>b</sup> | 9<sup>c</sup> | 1<sup>a</sup> | 4<sup>b</sup> | 7,5<sup>c</sup> | 6<sup>b</sup> | 2<sup>a</sup> | 8<sup>c</sup> | 2<sup>a</sup> | 2<sup>a</sup> | 11,5<sup>c</sup> | 6<sup>b</sup> | 7<sup>c</sup> | 8<sup>c</sup> |
| **Strength\*\*\*** |  |  |  |  |  |  |  |  |  |  |  |  |  |  |  |  |
| *Hip (max = 20)\** | 15<sup>b</sup> | 16<sup>b</sup> | 19<sup>a</sup> | 18<sup>a</sup> | 20 | 20 | 16<sup>b</sup> | 18<sup>a</sup> | 15<sup>b</sup> | 16<sup>b</sup> | 16<sup>b</sup> | 15<sup>b</sup> | 20 | 19<sup>a</sup> | 15<sup>b</sup> | 14<sup>c</sup> |
| *Knee (max = 10)\** | 7<sup>b</sup> | 7<sup>b</sup> | 8<sup>a</sup> | 8<sup>a</sup> | 10 | 8<sup>a</sup> | 8<sup>a</sup> | 8<sup>a</sup> | 6<sup>b</sup> | 8<sup>a</sup> | 8<sup>a</sup> | 7<sup>b</sup> | 9<sup>a</sup> | 10 | 7<sup>b</sup> | 7<sup>b</sup> |
| *Ankle (max = 25)\** | 17<sup>b</sup> | 20<sup>b</sup> | 25 | 20<sup>b</sup> | 25 | 21<sup>b</sup> | 13<sup>c</sup> | 20<sup>b</sup> | 11<sup>c</sup> | 19<sup>b</sup> | 17<sup>b</sup> | 13<sup>c</sup> | 12<sup>c</sup> | 20<sup>b</sup> | 15 | 11<sup>c</sup> |
| *Trunk (max = 10)\** | 7<sup>b</sup> |  | 10 |  | 10 |  | 9<sup>a</sup> |  | 10 |  | 7<sup>b</sup> |  | 7<sup>b</sup> |  | 6<sup>b</sup> |  |

*Measured values were categorized with <sup>a</sup> slight impairment, <sup>b</sup> considerable impairment and <sup>c</sup> severe* *impairment. No categorization means the value is within normal range.*
*\*values used for model personalization*
*\*\*sum of Modified Ashworth Scale*
*\*\*\*sum of different Muscle Testing Scores of muscles spanning the joint*
*/ no values recorded during the clinical examination.*

Soleus length was scaled based on maximal dorsiflexion angle with the knee flexed to 90°. The gastrocnemii length was scaled based on maximal dorsiflexion angle with the knees fully extended. Knee flexor length was scaled based on the bilateral popliteal angle. We used the same scaling factor for all knee flexors, i.e. biceps femoris long head, semimembranosus, semitendinosus, gracillis and sartorius. To obtain scaling factors for rectus femoris, we translated the score of the slow Duncan-Ely test to fixed scaling factors since one score allows for a wide range of knee flexion angles. For score 0 scaling was 1, for score 1 scaling was 0.9, and for score 2 scaling was 0.8 of nominal optimal fiber length.

We used a different approach to model iliopsoas contractures as their assessment is different. Iliopsoas contractures will lead to a different uni-(𝜃_*uni*_) and bilateral (𝜃_*bi*_) popliteal angle, that is the knee angle at maximal extension when the individual is supine with the thigh of the evaluated limb or both thighs vertical. When assessing the unilateral popliteal angle, the contralateral leg is laying down. Iliopsoas contractures will cause flexion of the contralateral hip and this will be compensated for by anterior pelvis tilt, which in turn will lead to increased hip flexion to position the thigh of the evaluated leg vertically. Increased hip flexion will in turn increase bi-articular hamstrings length and will thus lead to a larger knee extension deficit. increase in hip flexion angle based on the difference in popliteal angles and the ratio of the average moment arms of all bi-articular hamstrings with respect to the knee and hip:

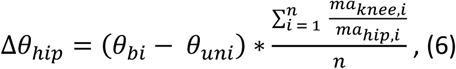

with *n* the number of bi-articular hamstrings, and *ma*_*knee,i*_ and *ma*_*hip,i*_ the moment arm of muscle *i* around knee and hip when in the position of the bilateral popliteal angle. We then determined the contracture of the contralateral iliopsoas by solving for the scaling factor that led to passive torque when the contralateral hip was extended beyond Δ𝜃_*hip*_.

Observed knee extension and plantar flexion deficits were modeled by shifting the coordinate limit torques that model the stiffness of the non-muscle soft tissues around the joint. For the knee, we shifted the limits such that torque started to develop at the observed angle for knee extension minus two degrees to obtain around 15 Nm torque at end range of motion. We modeled plantar flexion deficits based on the reported range of motion for the ankle towards plantar flexion. The onset of the coordinate limit torques was changed to 25° plantar flexion or 0° when the score was respectively ‘discrete’ or ‘severe’.

#### Modeling bony deformities

Bony deformities were modeled based on the MR images. First, bones were segmented with Materialise Mimics (Materialise, Leuven, BE). Further processing was performed in MuscleSegmenter [12]. We determined the pelvic frame based on anatomical landmarks of the pelvis and located the glycerin markers. Next, the hip joint center was estimated by fitting a sphere to the femoral head.

Then, reference frames of the femur, patella and tibia were determined based on anatomical landmarks. Next, the knee joint axis was determined based on fitting two ellipsoids to the condyles and visual inspection (congruence of joint contact surfaces) of the knee motion. Next, the paths of muscles spanning the hip and knee as well as the attachment points of the gastrocnemii on the femur were segmented. Muscle paths were chosen such that they approximated the centroid of the muscle belly. Next, the scaled generic model was updated by replacing the original joint centers, marker locations, and muscle paths by the values obtained from the MR images followed by linear scaling of optimal fiber lengths and tendon slack lengths (similar to the OpenSim Scale Tool) through a custom-made MATLAB (R2021b, MathWorks Inc, Natick, Massachusetts, USA) script.

Strength deficits observed during MMT can be due to both moment arm deficits or reduced muscle force generating capacity. The MRI-informed models already capture weakness due to moment arm deficits. Therefore, we modified the weakness scale factor derived from MMT to correct for weakness due to moment arm deficits:

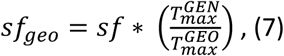

with *sf*_*geo*_ the corrected weakness scale factor to be used in the model with bony deformities, *sf* the weakness scale factor derived from MMT, 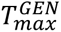 the maximal torque the muscle group could generate in the GEN model, and 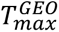 the maximal torque the muscle group could generate in the GEO model. *T*_*max*_ was evaluated with the model in the same position as during MMT and muscles activated to 100%.

Since muscle-tendon paths differ in the GEO model compared to GEN, we estimated contractures in both models separately.

### 3.5 Outcome measures

Experimental kinematics were calculated from marker trajectories collected during walking at self-selected speed based on the GEO model with OpenSim’s Inverse Kinematics Tool. Means and standard deviations of the experimental gait were calculated based on five to ten strides per side. Walking speed was determined based on the average forward speed of the posterior pelvis marker over at least four strides.

Root Mean Square Differences (RMSD) and Pearson correlations (r) were calculated between simulated and mean experimental kinematics for hip flexion, hip adduction, hip rotation, knee flexion and ankle dorsiflexion. We calculated the mean RMSD and correlation over all sagittal and non-sagittal plane degrees of freedom for each subject-model pair. Differences in median and interquartile range (IQR) between GEN and the other models were used to interpret the contribution of the modeled impairments to the altered gait. The minimal important difference for RMSD was defined to be 1.6° [43]. We deemed differences in correlation of 0.05 practically relevant for this study.

## 4. Results

### 4.1 Clinical Examination

The severity of impairments varied between individuals (Table 3). Common musculoskeletal impairments included contractures in the rectus femoris, hamstrings and ankle plantar flexors, and knee extension deficits. All participants had weakness but to a variable extent. Some participants had increased femoral anteversion and distal malalignments. Participants also had spasticity and reduced selectivity. The scale factors for optimal fiber length and active muscle force derived from the clinical assessment of muscle contractures and weakness are reported in the supplementary material (Table S5).

### 4.2 Predictive simulations

Overall, modeling impairments improved the agreement between simulated and measured hip flexion and knee flexion, but not hip adduction, hip rotation, and ankle dorsiflexion.

Differences between simulated kinematics based on GEN, representing how participants would walk in the absence of deficits except for segment length asymmetries, and measured kinematics varied greatly between participants (range RMSD sagittal: 7.2° - 25.5°) indicating large differences in kinematic deficits. Median RMSD and correlation between simulated kinematics based on GEN and measured kinematics were 12.8° (IQR 7.0°) and 0.82 (IQR 0.13) in the sagittal plane, and 8.4° (IQR 6.7°) and 0.23 (IQR 0.32) in the non-sagittal plane (Fig. 1). Median RMSD between simulated kinematics based on GEN and measured kinematics were 14.5° (IQR 12.0°) for hip flexion, 6.2° (IQR 4.8°) for hip adduction, 9.1° (IQR 8.5°) for hip rotation, 15.2° (IQR 8.4°) for knee flexion, and 6.0° (IQR 6.1°) for ankle dorsiflexion. Median correlation between simulated kinematics based on GEN and measured kinematics were 0.96 (IQR 0.09) for hip flexion, -0.18 (IQR 0.84) for hip adduction, 0.26 (IQR 0.58) for hip rotation, 0.92 (IQR 0.04) for knee flexion, and 0.62 (IQR 0.41) for ankle dorsiflexion.

**Fig. 1:**
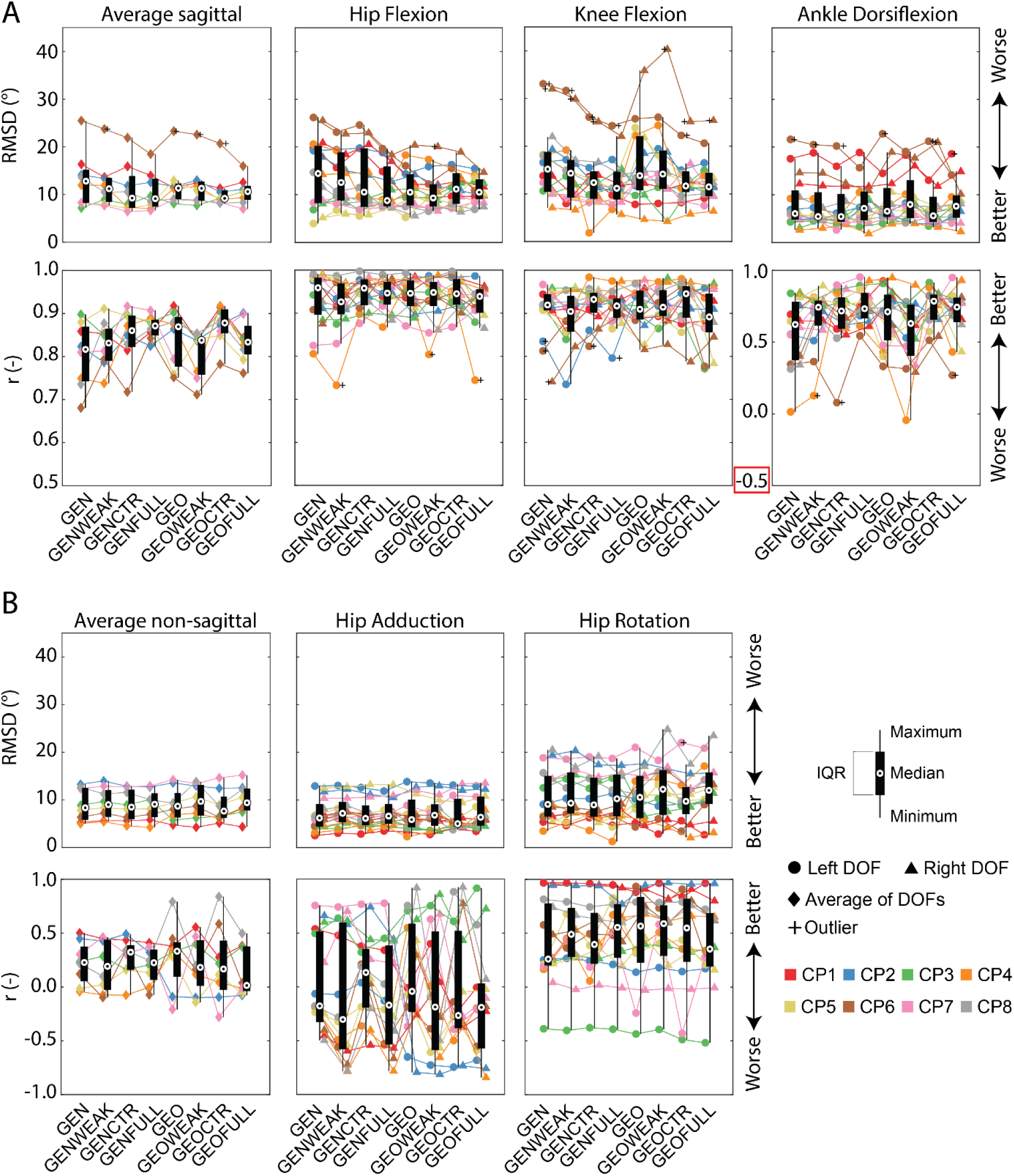
boxplots of average root mean square differences (RMSD) and Pearson correlation (r) for the eight models as well as results per degree of freedom (DOF) over the whole gait cycle. A) Sagittal plane DOFs. B) Non-sagittal plane DOFs. GEN is the generic scaled model, GENWEAK is GEN with weakness, GENCTR is GEN with contractures, GENFULL is GEN with both weakness and contractures. GEO is the model with MRI-based deformities, GEOWEAK is GEO with weakness, GEOCTR is GEO with contractures, and GEOFULL is GEO with both weakness and contractures. Average result per participant is the mean RMSD or correlation of sagittal or non-sagittal plane DOFs. Boxplots for the degrees of freedom are calculated based on the sixteen legs of the eight participants.

Modeling weakness (GENWEAK) mainly affected simulated hip kinematics (Fig. 1). In comparison to simulations based on GEN, modeling weakness reduced the RMSD between simulated and measured kinematics with 1.9° for hip flexion. The correlation increased with 0.12 for ankle dorsiflexion and with 0.23 for hip rotation, but also decreased with 0.13 for hip adduction (Table 4).

**Table 4.** Differences compared to the reference model GEN.

|  | GEN | GENWEAK | GENCTR | GENFULL | GEO | GEOWEAK | GEOCTR | GEOFULL |
| --- | --- | --- | --- | --- | --- | --- | --- | --- |
| **RMSD** |  |  |  |  |  |  |  |  |
| Average sagittal | 12.8° | **-1.6°** | **-3.5°** | **-3.7°** | -1.5° | -1.5° | **-3.6°** | **-2.2°** |
| Hip Flexion | 14.5° | **-1.9°** | **-3.9°** | **-5.7°** | **-3.9°** | **-5.1°** | **-3.3°** | **-3.9°** |
| Knee Flexion | 15.2° | -0.9° | **-2.8°** | **-4.0°** | -1.4° | -1.0° | **-3.5°** | **-3.7°** |
| Ankle Dorsiflexion | 6.0° | -0.6° | -0.6° | +1.1° | +0.5° | **+1.9°** | -0.4° | **+1.6°** |
| Average non-sagittal | 8.4° | +0.7° | +0.1° | +0.7° | +0.3° | +1.3° | -0.7° | +1.1° |
| Hip Adduction | 6.2° | +1.0° | -0.1° | +0.4° | -0.3° | +0.6° | -1.2° | +0.2° |
| Hip Rotation | 9.1° | +0.3° | -0.1° | +1.2° | +1.5° | **+3.2°** | +0.6° | **+2.9°** |
| **r** |  |  |  |  |  |  |  |  |
| Average sagittal | 0.87 | +0.01 | +0.04 | **+0.05** | **+0.05** | +0.02 | **+0.06** | +0.02 |
| Hip Flexion | 0.86 | -0.03 | +0.00 | -0.01 | -0.01 | -0.01 | -0.01 | -0.02 |
| Knee Flexion | 0.96 | -0.02 | +0.01 | +0.00 | -0.01 | -0.01 | +0.03 | -0.03 |
| Ankle Dorsiflexion | 0.89 | **+0.12** | **+0.09** | **+0.11** | **+0.09** | +0.01 | **+0.16** | **+0.12** |
| Average non-sagittal | 0.22 | -0.04 | **+0.10** | +0.00 | **+0.10** | **-0.05** | **-0.06** | **-0.21** |
| Hip Adduction | 0.04 | **-0.13** | **+0.31** | +0.01 | **+0.14** | -0.01 | **-0.09** | -0.01 |
| Hip Rotation | 0.77 | **+0.23** | **+0.14** | **+0.29** | **+0.31** | **+0.33** | **+0.29** | **+0.09** |

The root mean square difference (RMSD) and Pearson correlation (*r*) between predicted and experimental kinematics were compared between models. GEN is the generic scaled model, GENWEAK is GEN with weakness, GENCTR is GEN with contractures, GENFULL is GEN with both weakness and contractures. GEO is the model with MRI-based geometry, GEOWEAK is GEO with weakness, GEOCTR is GEO with contractures, and GEOFULL is GEO with both weakness and contractures. All models are compared to the reference model GEN. Values in bold are assumed to be practically relevant.

Modeling contractures (GENCTR) had the largest effect on sagittal plane hip and knee kinematics. In comparison to simulations based on GEN, modeling contractures reduced the RMSD between simulated and measured kinematics with 3.9° for hip flexion and 2.8° for knee flexion. Modeling contractures increased the correlation with 0.09 for ankle dorsiflexion, 0.31 for hip adduction, and 0.14 for hip rotation (Table 4).

Modeling contractures and weakness combined (GENFULL) resulted in the lowest average RMSD in the sagittal plane (Fig. 1). In comparison to simulations based on GEN the RMSD between simulated and measured kinematics was 5.7° lower for hip flexion and 4.0° lower for knee flexion. Modeling both weakness and contractures increased the correlation with 0.11 for ankle dorsiflexion and 0.29 for hip rotation (Table 4).

Modeling bony deformities (GEO) mainly affected simulated hip kinematics (Fig. 1). In comparison to simulations based on GEN, modeling bony deformities reduced the RMSD between simulated and measured kinematics with 3.9° for hip flexion and increased the correlation with 0.09 for ankle dorsiflexion, 0.14 for hip adduction and 0.31 for hip rotation (Table 4).

Modeling bony deformities and weakness combined (GEOWEAK) affected hip and ankle kinematics (Fig. 1). In comparison to simulations based on GEN, the RMSD between simulated and measured kinematics reduced with 5.1° for hip flexion but increased with 3.2° for hip rotation and 1.9° for ankle dorsiflexion. Modeling bony deformities and weakness increased the correlation with 0.33 for hip rotation (Table 4).

Modeling bony deformities and contractures combined (GEOCTR) affected the hip and knee kinematics (Fig. 1). In comparison to simulations based on GEN, the RMSD between simulated and measured kinematics reduced with 3.3° for hip flexion and 3.5° for knee flexion. Modeling bony deformities and contractures increased the correlation with 0.16 for ankle dorsiflexion and 0.29 for hip rotation, but decreased the correlation with 0.09 for hip adduction (Table 4).

Modeling all impairments combined (GEOFULL) did not result in the lowest RMSD or highest correlation (Fig. 1). In comparison to simulations based on GEN, the RMSD between simulated and measured kinematics reduced with 3.9° for hip flexion and 3.7° for knee flexion but increased with 2.9° for hip rotation. Modeling bony deformities, weakness and contractures together increased the correlation with 0.12 for ankle dorsiflexion and 0.09 for hip rotation (Table 4).

Gait deficits varied largely between participants as reflected in the large inter quartile range for both RMSD in the sagittal (7.0°) and non-sagittal plane (6.7°) between observed kinematics and simulated kinematics based on GEN (Fig. 1). The effect of modeling impairments varied largely between participants as reflected in the large range of changes in RMSD and correlation when modeling impairments (Fig. 1, Table 5, and supplement Fig. S5-S8). With model personalization (GEOFULL vs GEN), agreement between simulated and measured kinematics (based on RMSD and the correlation) improved for five out of eight participants, was mostly unaltered for one out of eight participants, and worsened for two out of eight participants in the sagittal plane (Table 5). In the non-sagittal plane, agreement between simulated and measured kinematics (based on RMSD and the correlation) improved for two out of eight participants, and was worse for six out of eight participants (Table 6).

**Table 5.** Differences between models per participant in the sagittal plane.

|  | CP1 | CP2 | CP3 | CP4 | CP5 | CP6 | CP7 | CP8 |
| --- | --- | --- | --- | --- | --- | --- | --- | --- |
| **RMSD** |  |  |  |  |  |  |  |  |
| GEN | 16.4° | 14.1° | 7.2° | 12.0° | 8.2° | 25.5° | 8.3° | 13.7° |
| GENWEAK | **-2.7°** | -0.8° | **+1.8°** | +0.0° | -0.3° | **-1.8°** | -0.6° | **-3.1°** |
| GENCTR | -0.4° | **-2.3°** | +0.8° | **-4.6°** | -1.1° | **-3.6°** | **-1.9°** | **-3.1°** |
| GENFULL | **-2.4°** | **-1.6°** | **+2.5°** | **-4.9°** | -0.4° | **-7.0°** | **-1.7°** | **-5.1°** |
| GEO | **-4.9°** | **-2.5°** | +0.9° | -0.7° | **+4.8°** | **-2.2°** | +0.6° | **-4.6°** |
| GEOWEAK | **-3.9°** | -1.1° | +0.5° | -0.2° | **+2.7°** | **-2.9°** | +0.4° | **-4.8°** |
| GEOCTR | **-5.0°** | **-3.6°** | **+2.0°** | **-3.0°** | +1.0° | **-4.8°** | +0.0° | **-5.5°** |
| GEOFULL | **-3.8°** | **-3.0°** | **+3.4°** | **-2.6°** | **+2.4°** | **-9.5°** | -1.3° | **-5.3°** |
| **r** |  |  |  |  |  |  |  |  |
| GEN | 0.86 | 0.82 | 0.90 | 0.75 | 0.88 | 0.68 | 0.81 | 0.74 |
| GENWEAK | -0.01 | -0.04 | **-0.05** | -0.01 | +0.03 | **+0.11** | +0.00 | **+0.14** |
| GENCTR | -0.04 | +0.03 | +0.00 | **+0.12** | +0.01 | +0.04 | **+0.11** | **+0.09** |
| GENFULL | +0.00 | +0.00 | -0.02 | **+0.14** | +0.00 | **+0.16** | **+0.10** | **+0.13** |
| GEO | **+0.06** | +0.04 | +0.00 | **+0.13** | **-0.12** | **+0.07** | -0.02 | **+0.15** |
| GEOWEAK | -0.02 | +0.01 | **-0.06** | +0.02 | -0.04 | +0.03 | **-0.06** | **+0.12** |
| GEOCTR | **+0.06** | **+0.05** | +0.01 | **+0.17** | -0.03 | +0.10 | **+0.05** | **+0.15** |
| GEOFULL | -0.02 | **+0.08** | **-0.06** | **+0.08** | **-0.09** | **+0.08** | **+0.09** | **+0.08** |

*Average of hip flexion, knee flexion and ankle dorsiflexion root mean square difference (RMSD) and Pearson correlation (r) between predicted and experimental kinematics. Results are shown as relative to GEN, the generic scaled model. GENWEAK is GEN with weakness, GENCTR is GEN with contractures, GENFULL is GEN with both weakness and contractures. GEO is the model with MRI-based geometries, GEOWEAK is GEO with weakness, GEOCTR is GEO with contractures, and GEOFULL is GEO with both weakness and contractures. Values in bold are assumed to be practically relevant.*

**Table 6.** Differences between models per participant in the non-sagittal plane.

|  | CP1 | CP2 | CP3 | CP4 | CP5 | CP6 | CP7 | CP8 |
| --- | --- | --- | --- | --- | --- | --- | --- | --- |
| **RMSD** |  |  |  |  |  |  |  |  |
| GEN | 5.3° | 13.3° | 8.9° | 5.0° | 7.8° | 6.4° | 12.4° | 12.7° |
| GENWEAK | +0.3° | +0.7° | +0.8° | +0.4° | +0.5° | +0.9° | +0.5° | -1.4° |
| GENCTR | +0.3° | -0.4° | +0.1° | -0.5° | +0.2° | +0.0° | +0.4° | -1.1° |
| GENFULL | +0.5° | -0.5° | +1.2° | -0.8° | +0.3° | +0.6° | +0.6° | **-2.6°** |
| GEO | -0.6° | +0.3° | **-3.1°** | **+1.8°** | +1.5° | **+1.7°** | **+1.8°** | **-3.3°** |
| GEOWEAK | -1.0° | -0.3° | **-2.6°** | **+2.2°** | **+2.8°** | **+2.3°** | +1.0° | +1.2° |
| GEOCTR | -0.2° | -0.8° | **-2.3°** | +1.1° | +1.1° | +0.0° | **+2.3°** | **-3.9°** |
| GEOFULL | -0.9° | -0.7° | -1.4° | **+3.2°** | **+2.7°** | **+2.0°** | **+2.8°** | -0.4° |
| **r** |  |  |  |  |  |  |  |  |
| GEN | 0.50 | 0.45 | 0.24 | -0.04 | -0.02 | 0.12 | 0.30 | 0.22 |
| GENWEAK | **-0.05** | -0.02 | **+0.05** | -0.03 | +0.04 | **-0.22** | **+0.16** | **-0.12** |
| GENCTR | **-0.07** | +0.04 | **-0.07** | -0.03 | **+0.17** | **+0.22** | +0.00 | **+0.12** |
| GENFULL | **-0.09** | **-0.13** | -0.03 | +0.04 | **+0.15** | **-0.19** | **+0.07** | +0.03 |
| GEO | **-0.12** | **-0.54** | **+0.09** | **+0.39** | **+0.30** | **+0.33** | **-0.51** | **+0.57** |
| GEOWEAK | **+0.05** | **-0.55** | **+0.16** | **+0.16** | **-0.13** | +0.04 | **+0.17** | -0.02 |
| GEOCTR | **-0.23** | **-0.54** | -0.02 | **+0.09** | **+0.13** | **+0.47** | **-0.58** | **+0.62** |
| GEOFULL | **-0.12** | **-0.53** | **+0.14** | +0.03 | **-0.05** | **-0.08** | **-0.31** | **+0.28** |

*Average of hip adduction and hip rotation root mean square difference (RMSD) and Pearson correlation (r) between predicted and experimental kinematics. Results are shown as relative to GEN, the generic scaled model. GENWEAK is GEN with weakness, GENCTR is GEN with contractures, GENFULL is GEN with both weakness and contractures. GEO is the model with MRI-based geometries, GEOWEAK is GEO with weakness, GEOCTR is GEO with contractures, and GEOFULL is GEO with both weakness and contractures. Values in bold are assumed to be practically relevant.*

Overall, the agreement between simulated and measured kinematics improved most when personalizing models for individuals whose gait is less well-represented by the generic model.

### 4.3 Exemplar individuals

We selected two exemplar individuals for a more in depth discussion but kinematic trajectories of all participants can be found in the supplement (Fig. S1-S4). CP3 had only limited gait deficits consisting mainly of hyperextension of the right knee during stance. In contrast, CP6 had considerable gait deficits reflected in the large RMSD between simulated kinematics based on GEN and measured kinematics (25.5° in the sagittal plane and 6.4° in the non-sagittal plane) and walked in crouch characterized by increased hip flexion, knee flexion and ankle dorsiflexion.

As expected given the mild deficits, simulations based on GEN closely matched the experimental gait kinematics in CP3. Modeling impairments did not consistently improve the agreement between simulated and measured kinematics. GEOWEAK was the only model able to replicate the extended knee throughout stance but at the cost of larger differences between simulated and measured ankle dorsiflexion.

Modeling impairments improved the agreement between simulated and measured kinematics for CP6 with simulations based on GEOFULL most closely matching experimental kinematics (Table 5). Nevertheless, differences between simulated and measured kinematics remained large even for the most personalized model (GEOFULL). Contractures contributed more to the crouch gait pattern than weakness or bony deformities (purple versus red/orange lines in Fig. 2).

**Fig. 2:**
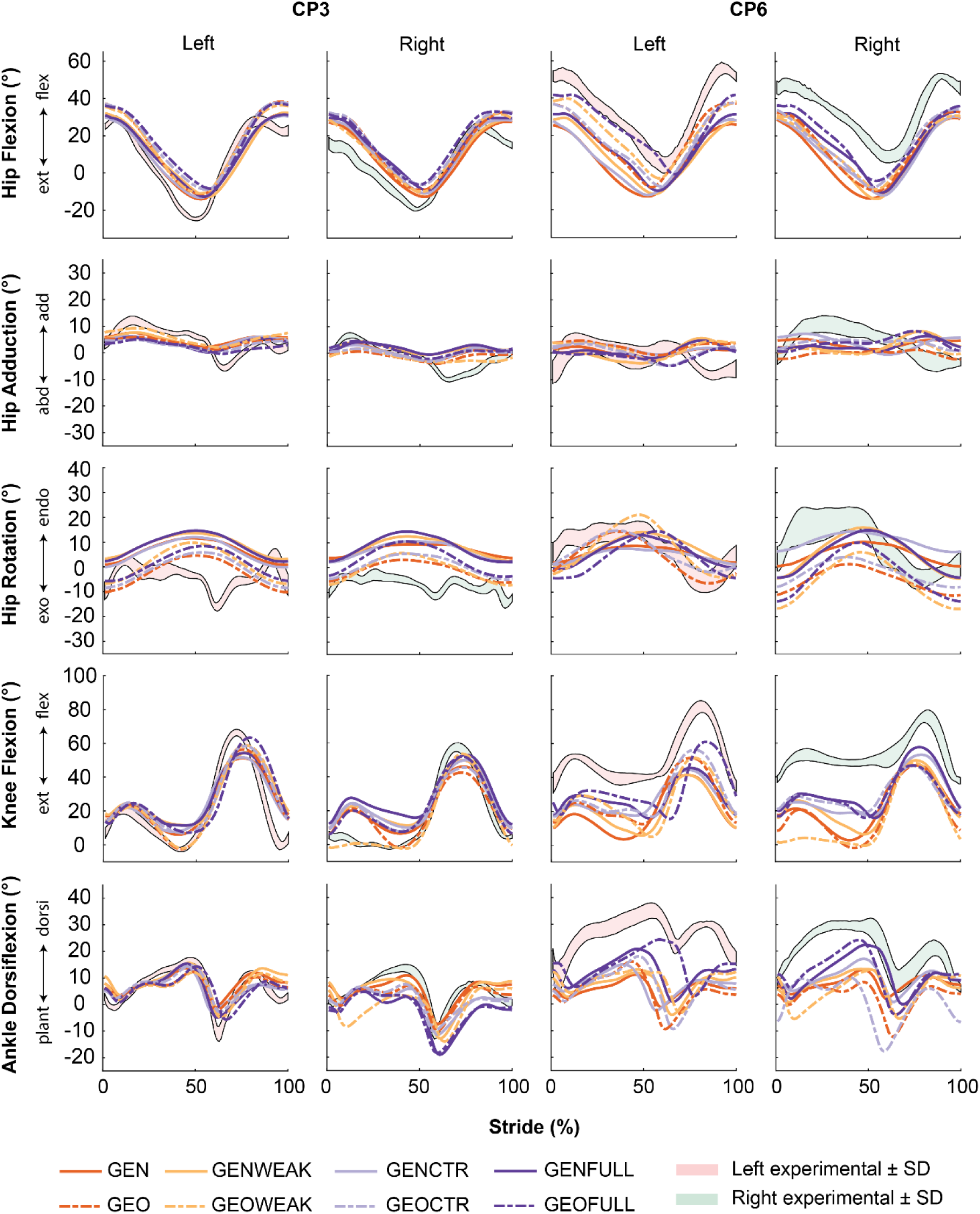
Experimental and simulated kinematics of two exemplar participants (CP3 and CP6). GEN is the generic scaled model, GENWEAK is GEN with weakness, GENCTR is GEN with contractures, GENFULL is GEN with both weakness and contractures. GEO is the model with MRI-based geometry, GEOWEAK is GEO with weakness, GEOCTR is GEO with contractures, and GEOFULL is GEO with both weakness and contractures. GEN represents how a typically developing individual with the same dimensions as the participant would walk. CP3 is representative for individuals with small kinematic deficits for whom modeled impairments do not explain kinematic deficits whereas CP6 is representative for individuals with large kinematic deficits for whom modeled impairments explain part of the kinematic deficits.

## 5. Discussion

Across the individuals with CP planned for MLS included in this study, modeled musculoskeletal impairments accounted for little to up to 39% of the observed gait deviations (RMSD, GEOFULL, CP8). Contractures had the largest effect on simulated kinematics when modeled in isolation, larger than either weakness or bony deformities in isolation. Interestingly, when combined, the contribution of all impairments was smaller than the sum of their isolated effects. The extent to which modeled impairments explained gait deficits was larger in the sagittal than in the non-sagittal plane and varied largely across individuals. However, between-subject variability in the differences between simulated and measured kinematics was reduced when modeling bony deformities. In addition, modeled impairments explained more of the gait deficits for more involved individuals. There are two possible reasons for the rather small contribution of the modeled impairments to gait deficits. First, motor control impairments and unmodeled deformities of tibia and foot might significantly contribute to gait deficits. Second, the accuracy of our modeling approach based on clinical data might be limited. Yet it is hard to distinguish these reasons based on our results. Overall, the limited and variable contribution of musculoskeletal deficits is in line with the limited and variable outcome of multilevel surgery targeting musculoskeletal deficits [44].

Our approach enables systematic assessment of the effect of both isolated impairments and their interactions. Overall, the largest effect was observed on sagittal plane kinematics (Fig. 1; decrease in sagittal plane RMSD up to 3.7° and decrease in non-sagittal plane RMSD only up to 0.7°), which is surprising since the modeled impairments were not limited to the sagittal plane. In the sagittal plane, contractures had the largest effect on the kinematics, but modeling bony deformities reduced the variability in RMSD suggesting that bony deformities explain some of the between-subject variability in gait deficits. Modeling impairments mostly affected hip kinematics. For the knee, only modeling contractures resulted in a closer match with the observed gait suggesting weakness and bony deformities were not the main contributors to observed deviations in knee kinematics. For ankle dorsiflexion, the correlation between simulated and experimental kinematics improved in all models except in the model with bony deformities and weakness, which was also the only model for which the RMSD between simulated and experimental kinematics increased with respect to GEN. In the non-sagittal plane, there were no major changes in RMSD with exception for hip rotation when modeling bony deformities and weakness. Contrary, the correlation between simulated and experimental hip rotations improved consistently when modeling impairments, either in isolation or in combination. Correlations between simulated and experimental hip abduction were low suggesting that the hip abduction pattern was not well captured across models. Although some care might be required when interpreting correlations for non-sagittal plane kinematics as we observed large within-subject trial-to-trial variability in the experimental data for these degrees of freedom.

The contribution of combined impairments was smaller than the sum of the contribution of isolated impairments, which illustrates the non-linear relationship between musculoskeletal impairments and gait kinematics. While both contractures and weakness in isolation affected gait kinematics, the magnitude of the change in kinematics (RMSD) due to their combined effect was similar to the effect of contractures in isolation. Hence, contractures might counteract some of the effects of weakness. On average, modeling bony deformities explained some of the gait deficits. Yet, the average effect of bony deformities, contractures, and weakness (GEOFULL) was smaller than the average effect of weakness and contractures (GENFULL). However, modeling bony deformities reduced the between-subject variability in RMSD between simulated and experimental kinematics. It is possible that we captured the variability in bony deformities, but that we introduced a systematic error since GEO was not constructed in the same way as GEN. GEN was constructed with muscle origin and insertion points defined relative to bone geometries and via points to avoid muscles intersecting with the bones [45] whereas the GEO model was constructed by manually determining muscle points such that the muscle paths were positioned in the center of the muscle volume. However, participants were laying in supine during the MRI-scan, which might have introduced changes in muscle shape with respect to muscle shape during walking and might therefore have led to inaccuracies in representing the muscle path. In addition, modeled bony deformities might interact with unmodeled tibia and foot deformities.

The contribution of modeled musculoskeletal impairments to altered gait kinematics varied largely between individuals. Modelling impairments yielded the largest reduction in the difference between simulated and measured kinematics when simulated kinematics based on GEN differed a lot from the experimental kinematics. The RMSD between measured kinematics and simulated kinematics based on GEN was below 8.4° for the three individuals (CP3, CP5, and CP7) for whom modeling impairments did not improve the agreement with experimental kinematics, whereas it was 12.0° to 25.5° for the individuals for whom modeling impairments did improve the agreement with experimental kinematics (Table 5). If our simulations are correct, this would suggest that correcting the modeled impairments, e.g. through MLS, would not improve gait kinematics in individuals with mild deviations in gait kinematics. This interpretation is in line with the finding from a data-driven study that MLS has a larger influence on the gait deviation index (GDI, a measure of deviations in gait kinematics) in children with a larger GDI pre-treatment with smaller gains or even declines in children with milder involvements [46]. However, we should be careful with interpreting these results until we can further validate our simulations. It is possible that the limited accuracy of our modeling and simulation approach explains why we only capture the effect of severe impairments as only severe impairments might cause alterations in kinematics that are larger than modeling errors. Yet, without modeling all impairments – including motor control impairments – it is hard to validate our simulations. We previously validated simulated kinematics in healthy individuals and found that RMSD normalized to the standard deviation between simulated and measured kinematics for hip, knee, and ankle angles were between 3.23° and 5.38° in stance and between 2.15° and 5.67° during swing with the smallest differences for hip adduction during swing and stance, and the largest differences for the ankle in stance and hip flexion in swing [33]. It was therefore unlikely that we would have obtained smaller RMSD here. In addition, we used geometric scaling to model musculoskeletal properties in children but body proportions and muscle properties are known to vary between adults and children [47]. In addition, optimal fiber length and tendon slack length were linearly scaled based on muscle-tendon length whereas linear scaling changes the operating range of the muscles especially when length differences are large as in some of the smaller children [48], [49]. In our model, 3D muscles were represented by line segments. This is especially challenging when defining muscle tendon paths based on MRI images, especially in children whose muscles are smaller and more difficult to identify.

The contribution of modeled impairments to gait deficits was limited suggesting an important contribution of unmodeled motor control impairments. Modeled impairments explained on average 17% and never more than 39% of the observed alterations in gait kinematics. This is largely in line with findings from another study that sought to unravel causal relationships between impairments and gait kinematics based on data-driven methods. Steele and Schwartz tested relationships between motor control and musculoskeletal impairments and the GDI [50]. The motor control and musculoskeletal impairments included in their data-driven analysis explained 63% of the variance in the pre-MLS GDI. They found that motor control impairments and strength (manual muscle testing scores) were most strongly related to the pre-SEMLS GDI, followed by knee extension and tibial torsion. We only studied the effect of muscle weakness, muscle contractures, and bony deformities, which might explain why our simulations captured less of the observed gait deviations. Contrary to their results, our simulations suggest that muscle contractures rather than muscle weakness have a higher contribution to altered gait. In addition, we showed that alterations in musculoskeletal geometry of hip and knee also contributed to alterations in gait kinematics, whereas Steele and Schwartz did not find a contribution for femoral anteversion. The results of Steele and Schwartz suggest that modeling tibial torsion is important for explaining alterations in gait kinematics, but we did not model tibial torsion here. It is likely that foot deformities, not considered in our study nor the study of Steele and Schwartz, also contribute to altered gait kinematics. We have previously shown that simulated gait kinematics are sensitive to properties of the foot including arch height and stiffness [33], [51], which are often altered in CP [52]. In this study, participants CP5 and CP6 had similar impairments based on the clinical examination but had very different gait patterns. Upon closer inspection of the movies collected during gait analysis, we observed that CP6 had a midfoot break, which might explain the large stance knee flexion not captured by the simulations. It is important to note that the contribution of musculoskeletal impairments might be larger when motor control impairments are considered as well, due to non-linear interactions between impairments [53].

While our approach for model personalization based on data from the clinical examination is attractive because it is broadly applicable, other methods might yield higher accuracy. The clinical examination is part of the standard clinical assessment of patients in many hospitals and therefore, personalization based on data from the clinical examination does not require additional assessments. Yet, manual muscle testing is based on a categorical strength scale and therefore we had to make assumptions on how the strength scales translated to a reduction in force output. In addition, the range of motion measures were not fully instrumented. The joint angles at end range of motion were measured with a goniometer, which is not the most accurate tool to measure joint angles [54], and the applied torque at end range of motion was not measured. We assumed that the torque at end range of motion was 15 Nm based on available data from instrumented assessments. However, although the clinical examinations were performed by trained and experienced clinicians, it is plausible that the actual torque deviated from 15 Nm. Alternative methods for estimating muscle parameters based on experimental data of muscle activity (through electromyography), joint kinematics, and joint moments across movements require additional data collection but may be more accurate because the input data is quantitative and contains more information, including time trajectories across passive and active movements [14], [15], [16], [17], [18], [19]. It remains to be investigated how sensitive the predicted gait kinematics are to the methods used for model personalization.

Finally, it is important to note that CP is a heterogeneous disease warranting caution when generalizing results. Here we included eight children and young adults with spastic CP planned for MLS spanning a wide age range, including both unilateral and bilateral CP, and presenting with different impairments and gait deviations. However, this does not represent the whole population of individuals with CP.

## 6. Conclusion

This study demonstrates the potential of predictive simulations based on subject-specific models to evaluate the impact of musculoskeletal impairments on gait in individuals with cerebral palsy. Our findings indicate that muscle weakness, contractures, and bony deformities contribute in a limited and variable manner, accounting for up to 39% of the observed gait deviations. These results suggest that other factors, such as motor control deficits and unmodeled skeletal abnormalities (e.g. tibial torsion and foot deformities) may play a significant role in shaping individual gait strategies. Additionally, limitations in the modeling process, particularly those related to assumptions and input derived from clinical examinations, may contribute to under- or overestimation of impairment effects. Future research should aim to enhance model fidelity by incorporating additional impairments and leveraging more quantitative experimental data. Doing so may improve the ability of simulations to inform treatment planning, particularly in complex and heterogeneous patient populations.

## 7. List of abbreviations

a: muscle activation
CP: Cerebral palsy
DOF: Degree of freedom
*Ė*: Metabolic energy rate
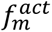: Active muscle force
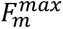: Maximal isometric force
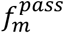: Passive muscle force
GDI: Gait deviation index
IQR: Interquartile range
*ma*: Moment arm
MLS: Multilevel surgery
MMT: Manual muscle testing
MRI: Magnetic resonance imaging
MSK: Musculoskeletal
RMSD: Root mean square difference
*sf*: scaling factor
𝜽_*bi*_: Bilateral popliteal angle
𝜽_*uni*_: Unilateral popliteal angle
*T*_*max*_: Maximal torque
*T*_*p*_: Passive joint torques
*u*_*a*_: Joint accelerations

## 8. Supplementary information

See Supplementary_material_v2-0-0.docx

## 8.1 Acknowledgements

The authors wish to thank all the children and parents for their participation in this study. We also thank the colleagues of the University Hospital Leuven who supported recruitment and data collection.

## 8.2 Author contributions

The study was conceptualized and designed by BVDB and FDG. Data collection was performed by BVDB. Software implementation was handled by BVDB, LD, and FDG. Modeling and simulations were conducted by BVDB. Data analysis and visualizations were carried out by BVDB and FDG. BVDB, LD, IJ, KD, AVC and FDG interpreted the results. The original draft was written by BVDB and FDG, while LD, IJ, KD, and AVC reviewed and edited the manuscript.

## 8.3 Funding

This research was funded by KU Leuven (C24M/19/064 awarded to F. De Groote) and Research Foundation Flanders (G0B4222N awarded to F. De Groote, 1SF1824N awarded to B. Van Den Bosch). The funders had no role in study design, data collection and analysis, decision to publish, or preparation of the manuscript.

## 8.4 Data availability

All data concerning this study is available within the manuscript and supplementary material. Detailed data and models are available upon reasonable request to the first author.

## 9. Declarations

### 9.1 Ethics approval and consent to participate

Data collection was approved by the local ethics committee (Ethical Committee UZ Leuven/KU Leuven; S64909) under the Declaration of Helsinki. The parents or participants’ caregivers provided written informed consent. Participants aged 12 years or older provided informed assent. All methodology adhered to the relevant regulations and guidelines.

### 9.2 Consent for publication

Not applicable.

### 9.3 Competing interests

The authors declare no competing interests.

## Notes

### Competing Interest Statement

The authors have declared no competing interest.

