## Supplementary material for "Estimating the contribution of musculoskeletal impairments to altered gait kinematics in children with cerebral palsy using predictive simulations"

1. Marker protocol

**Table S1**

*Marker protocol*

| **Segment** | **Marker ID** | **Anatomical reference position** | **Details** | **3D movement analyses** | **Glycerin markers MRI** |
| --- | --- | --- | --- | --- | --- |
| **Torso** | LSHO  RSHO | Left/Right acromion | Most proximal point. | x |  |
|  | CLAV | Upper edge of sternum/Jugular notch | In the middle. | x |  |
|  | STRN | Lower edge of sternum/Xiphoid process | M In the middle. | x |  |
|  | C7 | 7^th^ Cervical vertebrae | Spinous process.  In the middle. | x |  |
|  | T10 | 10^th^ Thoracic vertebrae | Spinous process.  In the middle. | x |  |
| **Pelvis** | LASI  RASI | Left/Right anterior superior iliac spine | Most pronounced part. | x | x |
|  | LPSI  RPSI | Left/Right posterior superior iliac spine | Below the dimple (if visible) or most pronounced part. | x | x |
|  | LLAT  RLAT | Left/Right lateral midpoint | Highest point of the iliac crest. | x | x |
| **Thigh** | LTHI  RTHI | Wand, Lower 1/3 of thigh (lateral side) |  | x |  |
|  | LF1  RF1 | Cluster marker thigh  Downward | Exact placement not critical. | x | x |
|  | LF2  RF2 | Cluster marker thigh  Dorsal | Exact placement not critical. | x | x |
|  | LF3  RF3 | Cluster marker thigh  Ventral | Exact placement not critical. | x | x |
|  | LKNE  RKNE | Lateral epicondyle of knee | Most pronounced part of epicondyle. Palpate from proximal with knee straight. Markers should hardly move while bending the knee. | x | x |
|  | LMEK  RMEK | Medial epicondyle of knee | Most pronounced part of epicondyle. Palpate from proximal with knee straight. Markers should hardly move while bending the knee | x | x |
| **Shank** | LTIB  RTIB | Wand, lower 1/3 of shank (anterior side) |  | x |  |
|  | LT1  RT1 | Cluster marker thigh Downward | Exact placement not critical. | x | x |
|  | LT2  RT2 | Cluster marker thigh  Dorsal | Exact placement not critical. | x | x |
|  | LT3  RT3 | Cluster marker thigh  Ventral | Exact placement not critical. | x | x |
| **Ankle** | LLANK  RLANK | Lateral malleolus | Most pronounced part. | x | x |
|  | LMANK  RMANK | Medial malleolus | Most pronounced part. | x | x |
|  | LSTL  RSTL | Sustaniculum Tali | Medial aspect of calcaneus, equidistant from HEE | x | x |
|  | LLCA  RLCA | Lateral calcaneus | Lateral aspect of calcaneus, equidistant from HEE | x | x |
| **Foot** | LHEE  RHEE | Calcaneus (dorsal part) | At same height as TOE, with the foot flat on the ground. | x | x |
|  | LTOE  RTOE | 2^nd^ Metatarsal head | Over the 2th metatarsal head on the midfoot side of the equines break between for- and midfoot. | x | x |
|  | LCM5  RCM5 | 5^th^ Metatarsal head | On top of 5^th^ metatarsal head. | x | x |
| Total amount of markers functional movements: 34 + 4 clusters of 3 = **46** | | | | | |
| Total amount of markers MRI: **36** | | | | | |

2. Ground contact model scaling

In all models, the radii ($r_{CS}$), stiffness ($k_{CS}$), and damping ($d_{CS}$) of the contact spheres were scaled to account for anthropometric differences (length and mass) between participants:

$r_{CS, subject}=r_{CS, DHondt{2024}_{3seg}}*\left( \frac{L_{subject}^{foot}}{L_{DHondt{2024}_{3seg}}^{foot}} \right)$ , (s1)

$k_{CS,subject}=k_{CS, DHondt{2024}_{3seg}}*\left( \frac{M_{subject}}{M_{DHondt{2024}_{3seg}}}* \left( \frac{L_{DHondt{2024}_{3seg}}}{L_{subject}} \right)^{2} \right)$ , (s2)

$d_{CS,subject}=d_{CS,DHondt{2024}_{3seg}}*\left( \frac{L_{subject}}{L_{DHondt{2024}_{3seg}}} \right)$ , (s3)

were $L^{foot}$ refers to the length of the foot, $L$ refers to body height and $M$ refers to body weight.

3. Manual Muscle Testing

The strength is evaluated for the full active range of motion (ROM). When the active range of motion is smaller than the passive range of motion (for instance due to co-contraction of the antagonists), the strength-score is not reduced, but the limits in active range of motion are noted, and the score for selectivity is reduced.

**Table S2**

*Manual Muscle Testing scoring system*

| **Score** | **Description** |
| --- | --- |
| 0 | Contraction cannot be palpated |
| 1 | Evidence of slight contraction of the muscle, but joint motion is not visible |
| 2 | Complete ROM in gravity eliminated plane (ROM can be slightly decreased because of co-contraction) |
| 3 | Perfect motion against gravity (almost full available ROM, ROM can be slightly decreased because of co-contraction) |
| 4 | Motion against gravity with some (moderate) resistance (full available ROM) |
| 5 | Motion against gravity with maximal resistance (full available ROM) |

In children with CP a lack of control of pelvis and trunk motion is frequently observed. Therefore, a specific evaluation of abdominal and back muscles is also performed.

**Table S3**

*Manual Muscle Testing scoring system for abdominal muscles*

| **Score** | **Description** |
| --- | --- |
| 0 | No visible / palpable abdominal contraction |
| 1 | Palpable contraction, but cannot elicit cervical and trunk flexion |
| 2 | Can raise head off the mat into cervical flexion |
| 3 | Can lift shoulders and scapulae off the mat |
| 4 | Can perform thoracic flexion |
| 5 | Can sit up with trunk flexion |

**Table S4**

*Manual Muscle Testing scoring system for back muscles*

| **Score** | **Description** |
| --- | --- |
| 0 | Cannot attempt any movement and no contraction can be palpated |
| 1 | Palpable contraction as the patient performs cervical extension (to raise head off the mat) |
| 2 | Can raise head and shoulders off the mat |
| 3 | Can raise chest and ribs off the mat |
| 4 | Can perform lumbar extension |
| 5 | Can perform lumbar extension with hip extension |

4. Scaling factors for modeling contractures and weakness

**Table S5**

*Scaling factors for modeling contractures and weakness*

|  | CP1 | | | CP2 | | CP3 | | | | CP4 | | CP5 | | | CP6 | | | CP7 | | | | CP8 | |
| --- | --- | --- | --- | --- | --- | --- | --- | --- | --- | --- | --- | --- | --- | --- | --- | --- | --- | --- | --- | --- | --- | --- | --- |
|  | L | R | L | | R | | L | R | L | | R | | L | R | | L | R | | L | R | L | | R |
| Optimal fiber length (GEN) |  |  |  | |  | |  |  |  | |  | |  |  | |  |  | |  |  |  | |  |
| *Soleus* | - | - | 0.81 | | 0.67 | | 0.78 | 0.65 | 0.69 | | - | | - | - | | - | - | | 0.58 | 0.75 | - | | - |
| *Gastrocs* | - | - | - | | 0.90 | | 1.03 | - | 0.93 | | - | | - | - | | - | - | | 0.76 | 0.82 | - | | - |
| *Hamstrings* | 0.76 | 0.78 | 0.82 | | 0.83 | | 0.94 | 0.92 | 0.73 | | 0.83 | | 0.76 | 0.84 | | 0.81 | 0.76 | | 0.88 | 0.90 | 0.83 | | 0.80 |
| *Rectus femoris* | 0.80 | 0.80 | - | | 0.90 | | - | 0.90 | - | | 0.80 | | - | - | | - | - | | 0.90 | 0.90 | - | | - |
| *Iliopsoas* | - | - | - | | - | | - | - | 0.65 | | 0.90 | | - | - | | 0.78 | - | | 0.90 | 0.92 | 0.75 | | 0.76 |
| Optimal fiber length (GEO) |  |  |  | |  | |  |  |  | |  | |  |  | |  |  | |  |  |  | |  |
| *Soleus* | - | - | 0.86 | | 0.68 | | 0.80 | 0.67 | 0.79 | | - | | - | - | | - | - | | 0.61 | 0.80 | - | | - |
| *Gastrocs* | - | - | - | | 0.88 | | 1.04 | 1.04^a^ | 0.82 | | - | | - | - | | - | - | | 0.80 | 0.88 | - | | - |
| *Hamstrings* | 0.76 | 0.84 | 0.97 | | 0.94 | | 0.97 | 0.88 | 0.83 | | 0.95 | | 0.83 | 0.85 | | 0.81 | 0.76 | | 1.01 | 1.11 | 0.84 | | 0.80 |
| *Rectus femoris* | 0.80 | 0.80 | - | | 0.90 | | - | 0.90 | - | | 0.80 | | - | - | | - | - | | 0.90 | 0.90 | - | | - |
| *Iliopsoas* | - | - | - | | - | | - | - | 1.10 | | 1.35 | | - | - | | 0.94 | - | | 1.19 | 1.22 | 0.94 | | 0.93 |
| Strength |  |  |  | |  | |  |  |  | |  | |  |  | |  |  | |  |  |  | |  |
| *Hip abductors* | 0.30 | 0.50 | 0.70 | | 0.50 | | 0.70 | 0.70 | 0.50 | | 0.50 | | 0.30 | 0.50 | | 0.30 | 0.30 | | 0.70 | 0.70 | 0.50 | | 0.30 |
| *Hip flexors* | 0.50 | 0.50 | 0.70 | | 0.70 | | 0.70 | 0.70 | 0.50 | | 0.70 | | 0.50 | 0.50 | | 0.70 | 0.70 | | 0.70 | 0.70 | 0.50 | | 0.50 |
| *Hip extensors* | 0.50 | 0.50 | 0.50 | | 0.50 | | 0.70 | 0.70 | 0.50 | | 0.50 | | 0.30 | 0.30 | | 0.30 | 0.30 | | 0.70 | 0.50 | 0.30 | | 0.30 |
| *hip adductors* | 0.50 | 0.50 | 0.70 | | 0.70 | | 0.70 | 0.70 | 0.50 | | 0.70 | | 0.70 | 0.70 | | 0.70 | 0.50 | | 0.70 | 0.70 | 0.50 | | 0.50 |
| *Knee flexors* | 0.30 | 0.30 | 0.50 | | 0.50 | | 0.70 | 0.50 | 0.50 | | 0.50 | | 0.30 | 0.50 | | 0.30 | 0.30 | | 0.70 | 0.70 | 0.30 | | 0.30 |
| *Knee extensors* | 0.50 | 0.50 | 0.50 | | 0.50 | | 0.70 | 0.50 | 0.50 | | 0.50 | | 0.30 | 0.50 | | 0.70 | 0.50 | | 0.50 | 0.70 | 0.50 | | 0.50 |
| *Ankle plantarflexors* | 0.30 | 0.30 | 0.50 | | 0.30 | | 0.50 | 0.30 | 0.20 | | 0.30 | | 0.20 | 0.30 | | 0.20 | 0.20 | | 0.20^b^ | 0.30^b^ | 0.20 | | 0.20 |
| *Ankle inversors* | 0.30 | 0.50 | 0.70 | | 0.50 | | 0.70 | 0.50 | 0.20 | | 0.50 | | 0.30 | 0.50 | | 0.30 | 0.20 | | 0.30 | 0.70 | 0.30 | | 0.20 |
| *Ankle eversors* | 0.30 | 0.50 | 0.70 | | 0.50 | | 0.70 | 0.70 | 0.20 | | 0.50 | | 0.10 | 0.30 | | 0.30 | 0.20 | | 0.30 | 0.70 | 0.30 | | 0.20 |
| *Ankle dorsiflexors* | 0.30 | 0.50 | 0.70 | | 0.50 | | 0.70 | 0.50 | 0.30 | | 0.50 | | 0.20 | 0.50 | | 0.50 | 0.30 | | 0.30 | 0.70 | 0.30 | | 0.20 |
| *Abdominal muscles* | 0.30 | | | 0.70 | | 0.70 | | | | 0.50 | | 0.70 | | | 0.30 | | | 0.30 | | | | 0.30 | |
| *Back muscles* | 0.50 | | | 0.70 | | 0.70 | | | | 0.70 | | 0.70 | | | 0.50 | | | 0.50 | | | | 0.30 | |

*- the scaling factor is 1*

*^a^ due to same value clinical exam for 0° and 90°*

*^b^ no value recorded during clinical exam, so this is based on the value of the previous clinical exam of this patient*

5. Moment arm deficits

**Table S6**

|  | CP1 | CP2 | CP3 | CP4 | CP5 | CP6 | CP7 | CP8 |
| --- | --- | --- | --- | --- | --- | --- | --- | --- |
| Hip Abductors R | 0.70 | 0.69 | 0.77 | 0.81 | 0.81 | 0.67 | 0.83 | 0.82 |
| Hip Flexors R | 3.07 | 3.73 | 1.46 | 0.90 | 0.96 | 2.35 | 1.33 | 0.92 |
| Hip Extensors R | 0.71 | 0.86 | 0.69 | 0.76 | 0.73 | 0.55 | 0.77 | 0.65 |
| Hip Adductors R | 0.78 | 1.06 | 0.75 | 0.93 | 0.61 | 0.86 | 0.94 | 0.53 |
| Knee Flexors R | 0.84 | 1.42 | 1.54 | 0.92 | 1.02 | 1.17 | 1.40 | 0.92 |
| Knee Extensors R | 1.31 | 1.25 | 1.99 | 0.85 | 1.01 | 1.15 | 1.56 | 0.66 |
| Ankle Plantar Flexors R | 1.09 | 1.04 | 1.07 | 1.13 | 0.85 | 0.79 | 1.22 | 1.16 |
| Ankle Invertors R | 1.15 | 1.06 | 1.09 | 1.17 | 0.95 | 0.82 | 1.10 | 1.12 |
| Ankle Evertors R | 1.12 | 1.04 | 1.08 | 1.14 | 0.99 | 0.85 | 1.04 | 1.09 |
| Ankle Dorsiflexors R | 1.00 | 0.99 | 1.00 | 1.02 | 0.98 | 0.83 | 0.99 | 1.00 |
| Hip abductors L | 0.80 | 0.91 | 0.86 | 0.84 | 0.86 | 0.68 | 0.62 | 0.90 |
| Hip Flexors L | 1.10 | 1.78 | 1.27 | 1.82 | 1.18 | 0.91 | 1.12 | 1.49 |
| Hip Extensors L | 0.72 | 1.13 | 0.69 | 0.42 | 0.70 | 0.68 | 0.92 | 0.72 |
| Hip Adductors L | 0.84 | 0.97 | 0.86 | 0.89 | 0.85 | 0.73 | 0.91 | 0.58 |
| Knee Flexors L | 1.27 | 1.82 | 1.13 | 1.09 | 1.09 | 1.07 | 1.41 | 0.87 |
| Knee Extensors L | 1.34 | 1.39 | 1.17 | 1.10 | 1.02 | 0.96 | 1.22 | 0.49 |
| Ankle Plantar Flexors L | 1.12 | 1.06 | 1.05 | 1.12 | 0.82 | 0.80 | 1.17 | 1.12 |
| Ankle Invertors L | 1.16 | 1.08 | 1.07 | 1.16 | 0.93 | 0.84 | 1.06 | 1.06 |
| Ankle Evertors L | 1.13 | 1.07 | 1.06 | 1.14 | 0.98 | 0.88 | 1.01 | 1.04 |
| Ankle Dorsiflexors L | 1.01 | 0.97 | 1.00 | 1.01 | 0.97 | 0.85 | 0.97 | 0.97 |
| Abdominal muscles | 0.96 | 0.95 | 0.92 | 1.00 | 0.84 | 0.82 | 0.85 | 0.90 |
| Back muscles | 0.99 | 1.00 | 0.94 | 1.03 | 0.87 | 0.85 | 0.87 | 0.92 |

*Moment arm deficits*

*We estimated moment arm deficits based on the ratio of maximal torque per muscle group (i.e. muscle activation 1) of the GEN and GEO models in the posture assumed during manual muscle testing. Values > 1 represent larger moment arms in GEO, values < 1 represent smaller moment arms in GEO.*

6. Simulated and experimental kinematics per subject

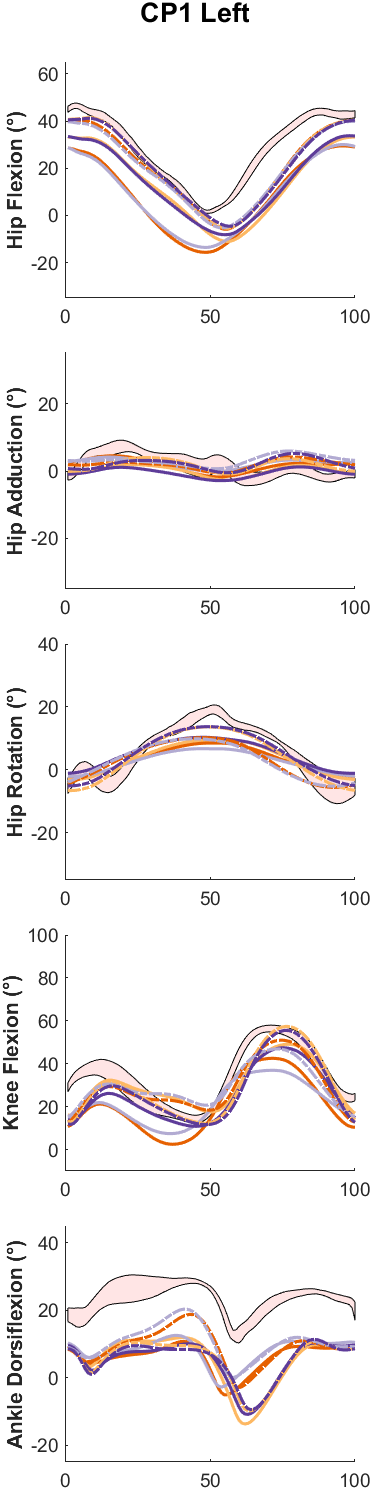

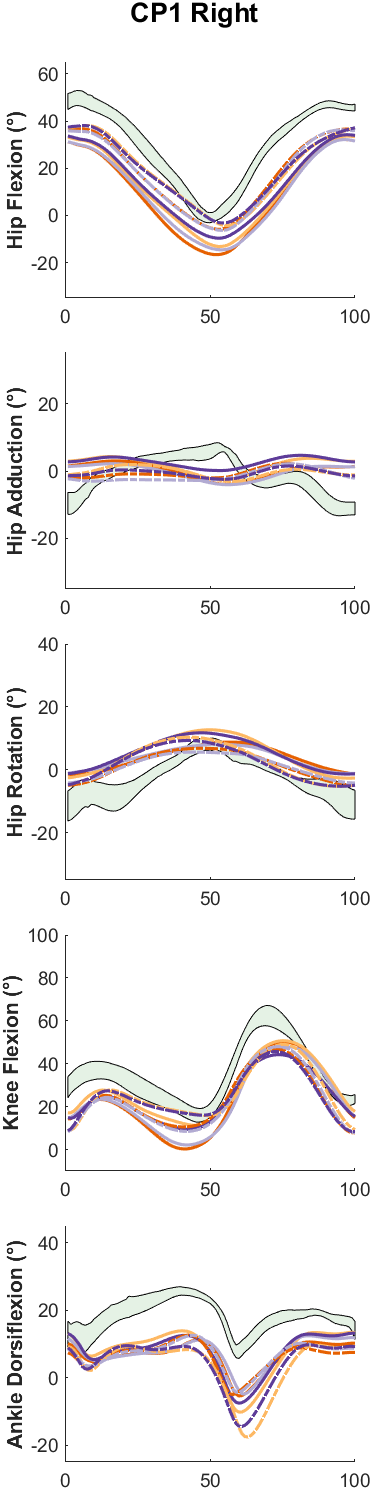

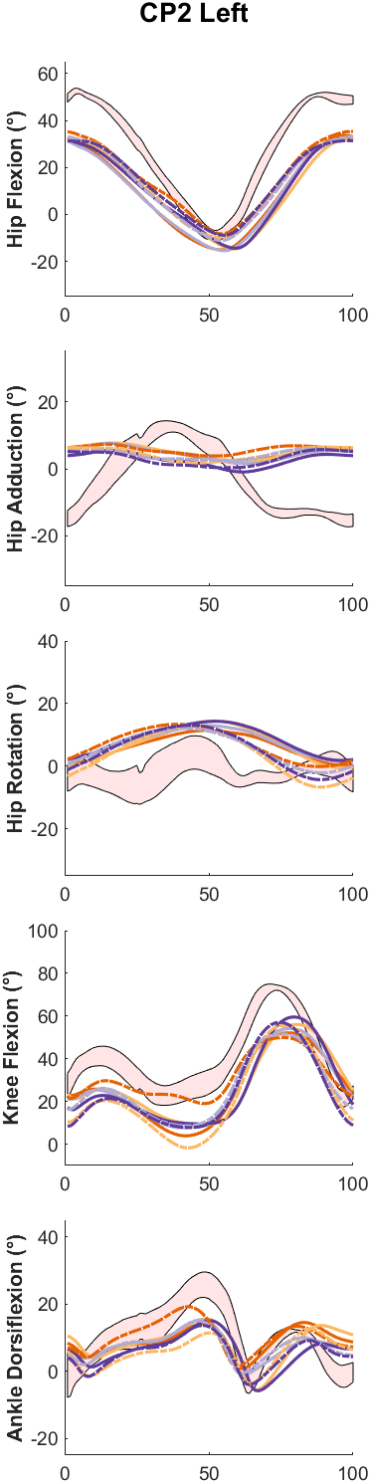

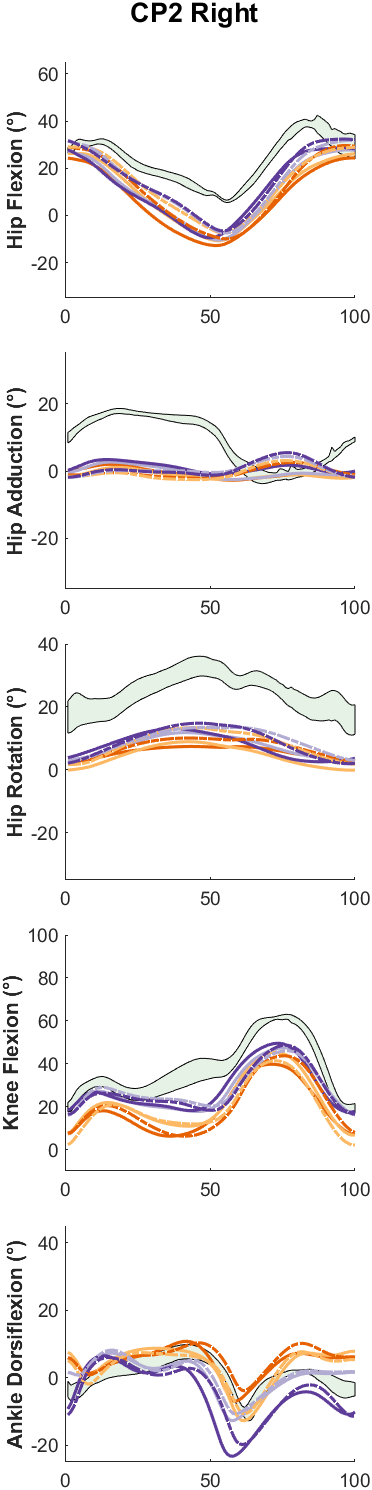

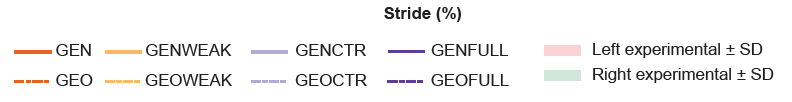

*Fig. S1:* *Experimental and simulated kinematics of CP1 and CP2. GEN is the generic scaled model, GENWEAK is GEN with weakness, GENCTR is GEN with contractures, GENFULL is GEN with both weakness and contractures. GEO is the model with MRI-based deformities, GEOWEAK is GEO with weakness, GEOCTR is GEO with contractures, and GEOFULL is GEO with both weakness and contractures. GEN represents how a typically developing individual with the same dimensions as the patient would walk.*

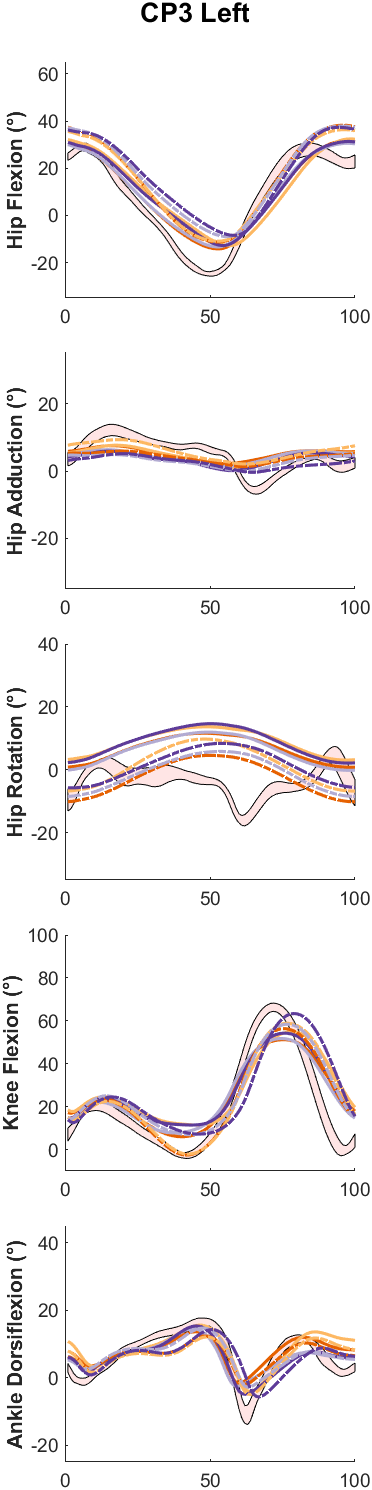

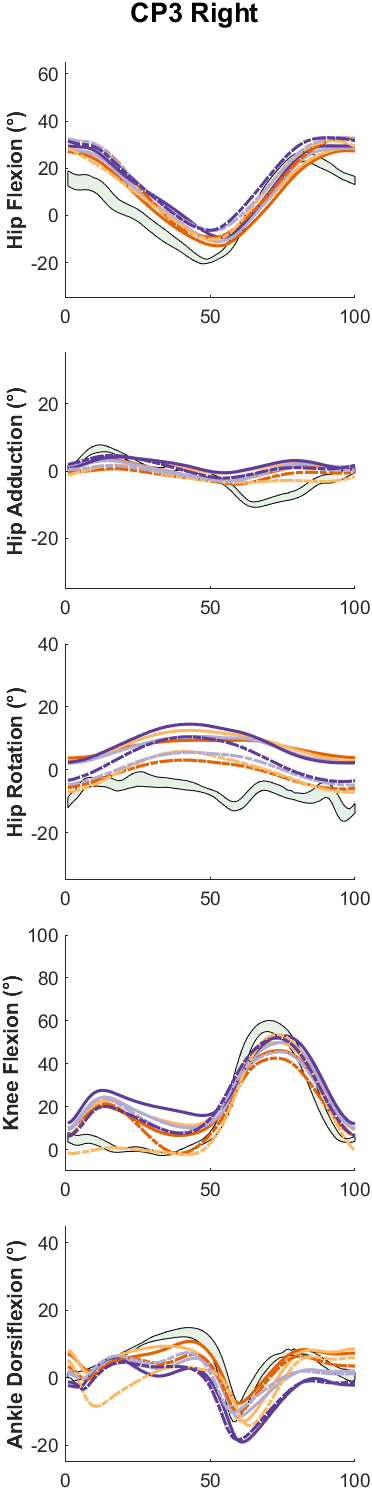

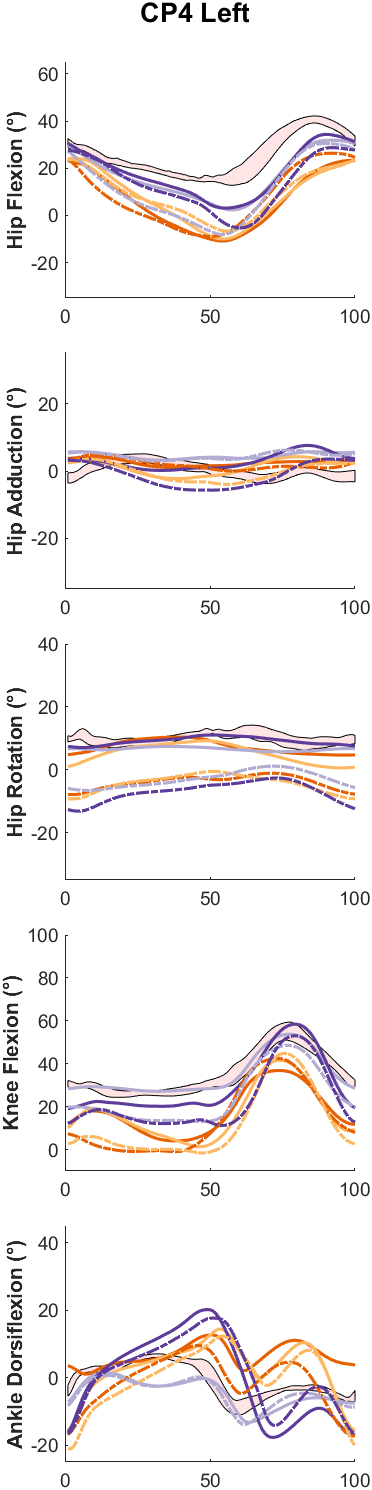

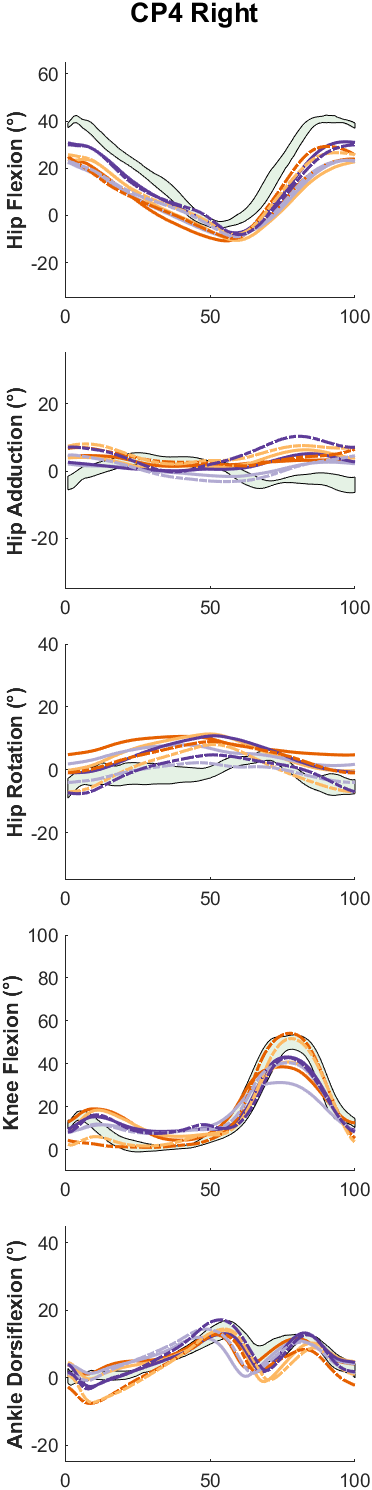

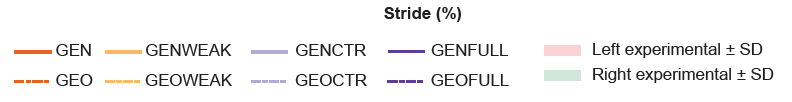

*Fig. S2:* *Experimental and simulated kinematics of CP3 and CP4. GEN is the generic scaled model, GENWEAK is GEN with weakness, GENCTR is GEN with contractures, GENFULL is GEN with both weakness and contractures. GEO is the model with MRI-based deformities, GEOWEAK is GEO with weakness, GEOCTR is GEO with contractures, and GEOFULL is GEO with both weakness and contractures. GEN represents how a typically developing individual with the same dimensions as the patient would walk.*

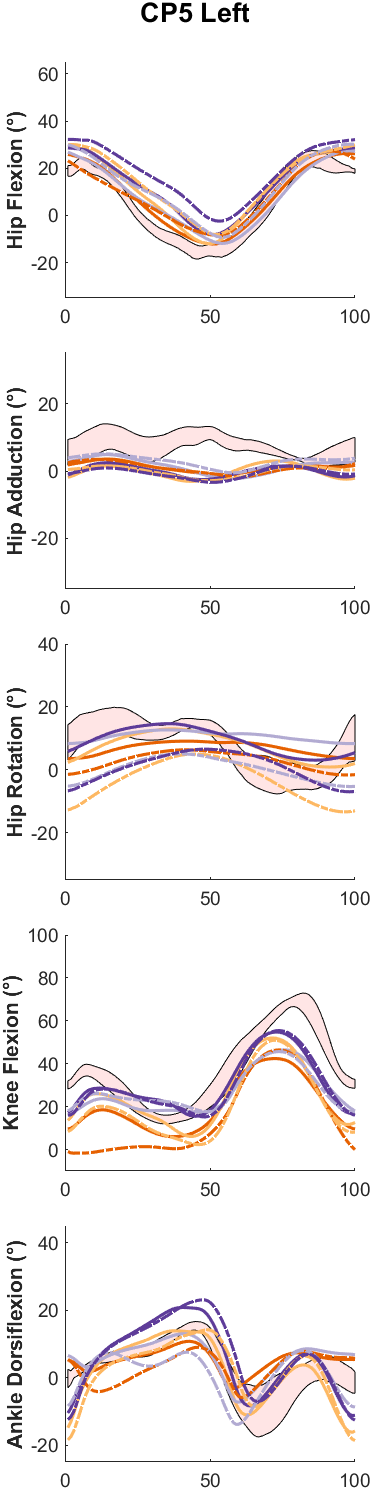

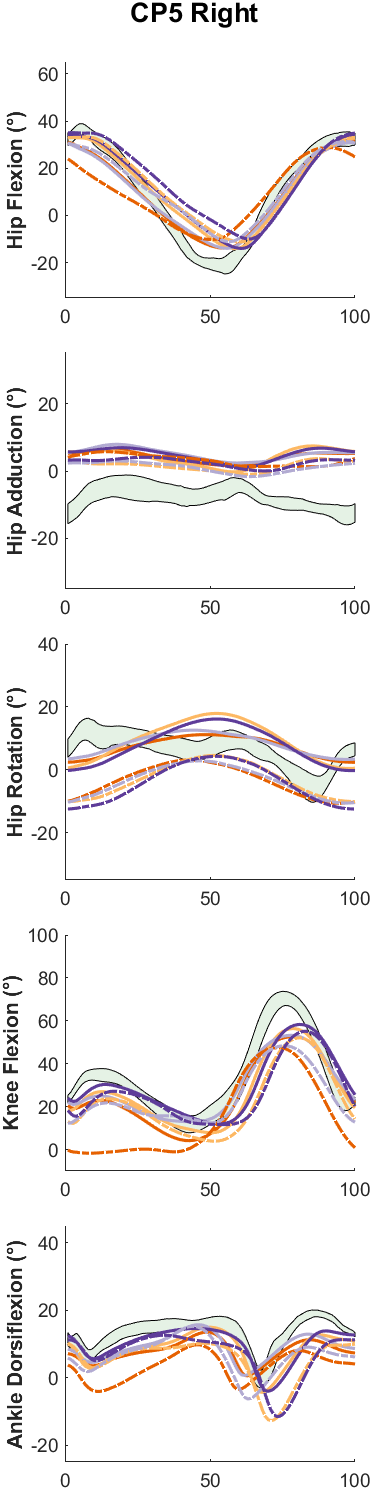

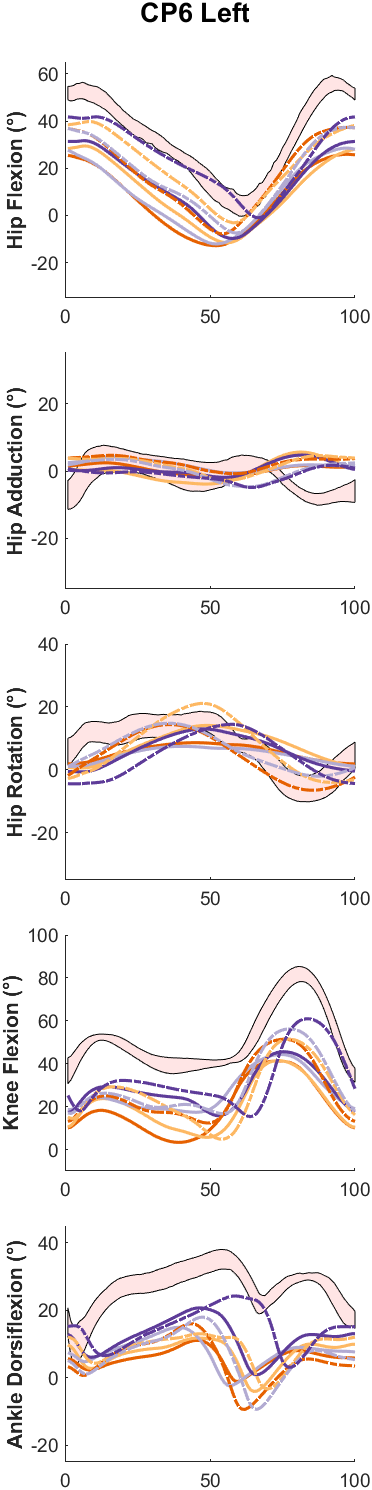

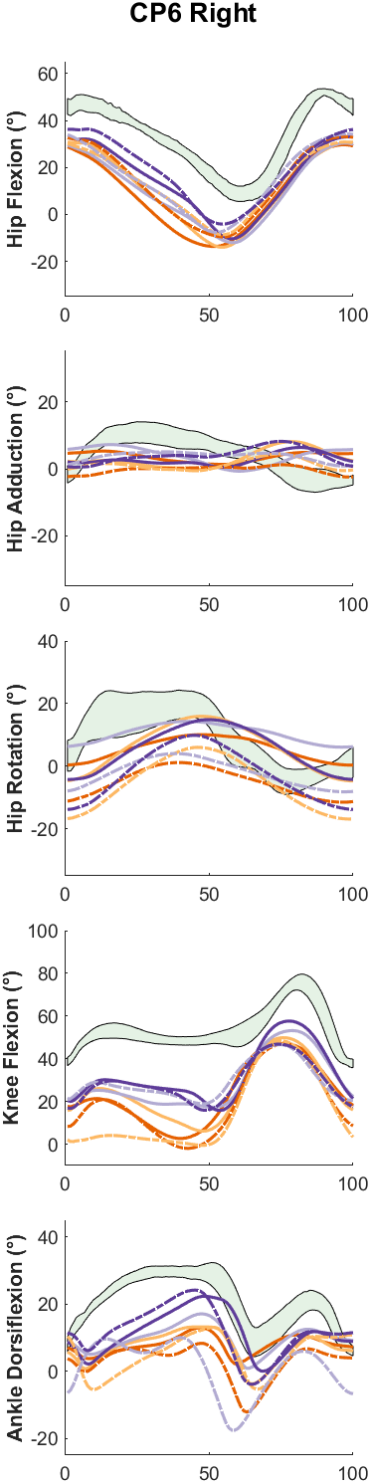

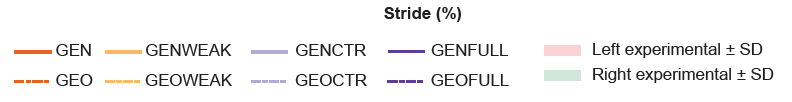

*Fig. S3:* *Experimental and simulated kinematics of CP5 and CP6. GEN is the generic scaled model, GENWEAK is GEN with weakness, GENCTR is GEN with contractures, GENFULL is GEN with both weakness and contractures. GEO is the model with MRI-based deformities, GEOWEAK is GEO with weakness, GEOCTR is GEO with contractures, and GEOFULL is GEO with both weakness and contractures. GEN represents how a typically developing individual with the same dimensions as the patient would walk.*

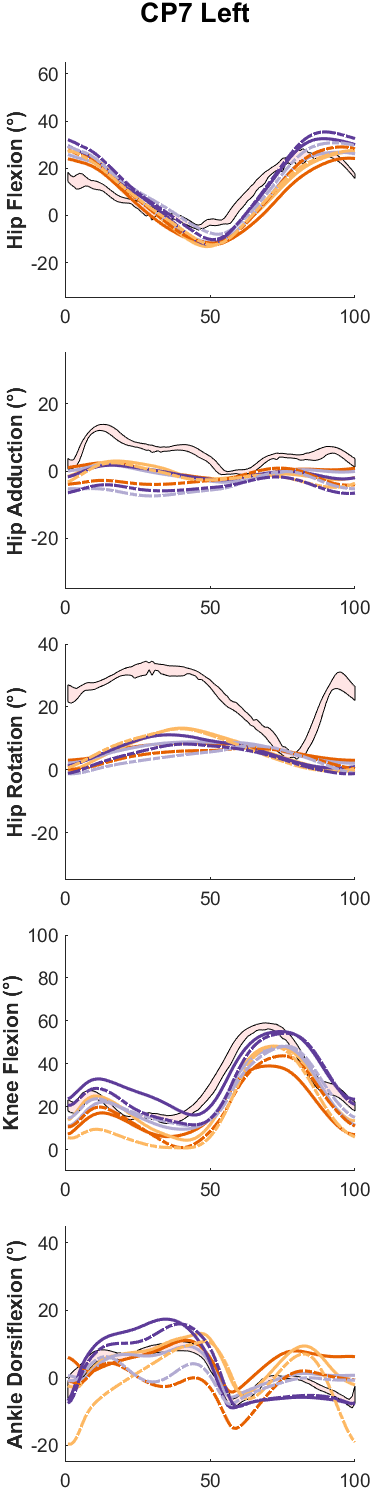

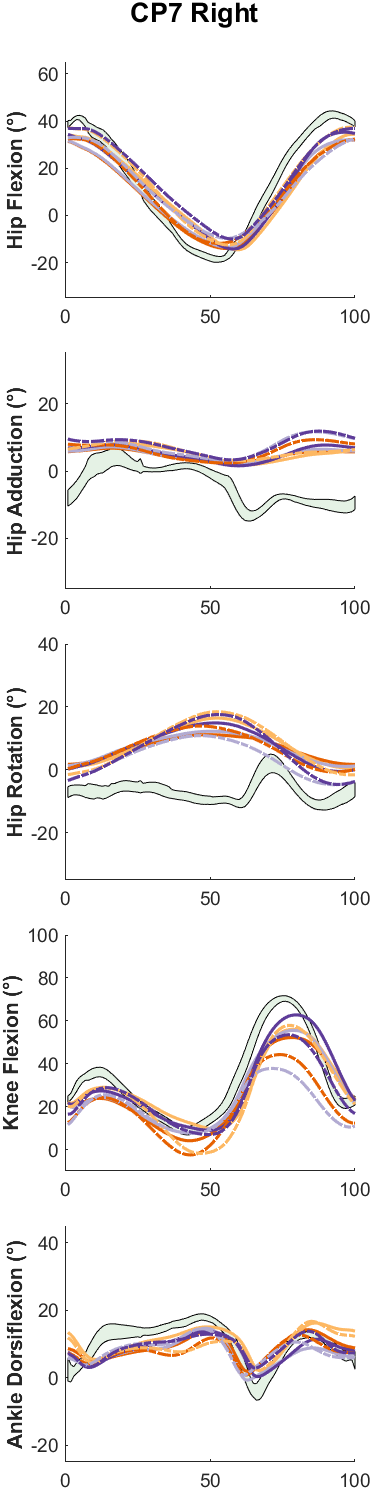

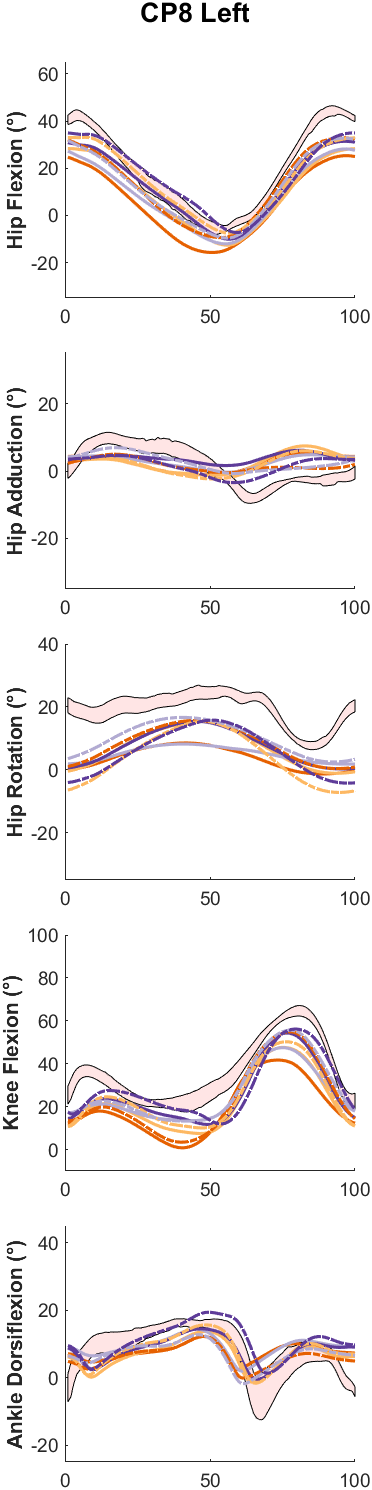

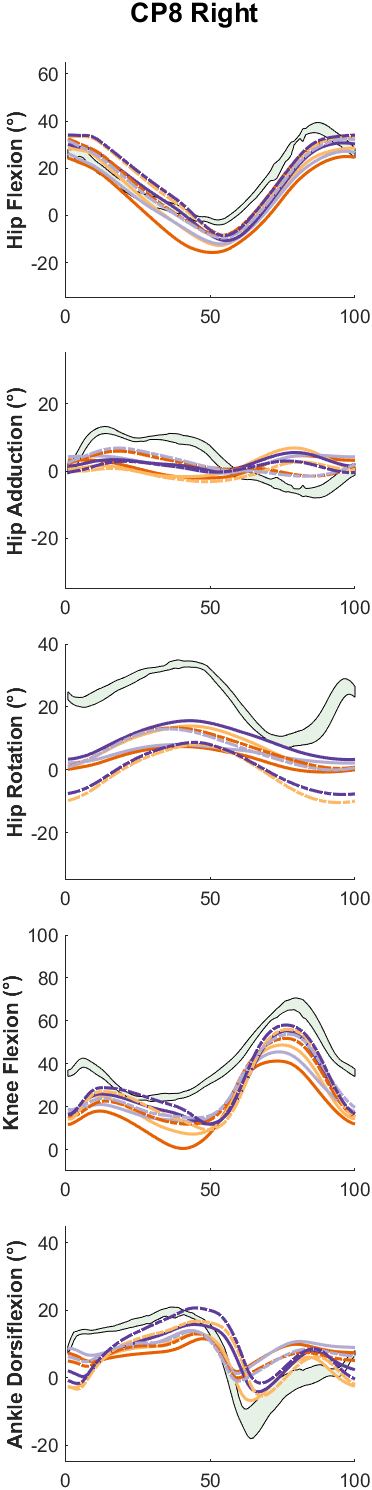

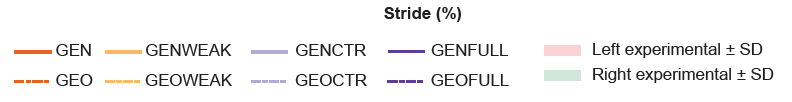

*Fig. S4:* *Experimental and simulated kinematics of CP7 and CP8. GEN is the generic scaled model, GENWEAK is GEN with weakness, GENCTR is GEN with contractures, GENFULL is GEN with both weakness and contractures. GEO is the model with MRI-based deformities, GEOWEAK is GEO with weakness, GEOCTR is GEO with contractures, and GEOFULL is GEO with both weakness and contractures. GEN represents how a typically developing individual with the same dimensions as the patient would walk.*

7. RMSD and CC differences between models, per subject

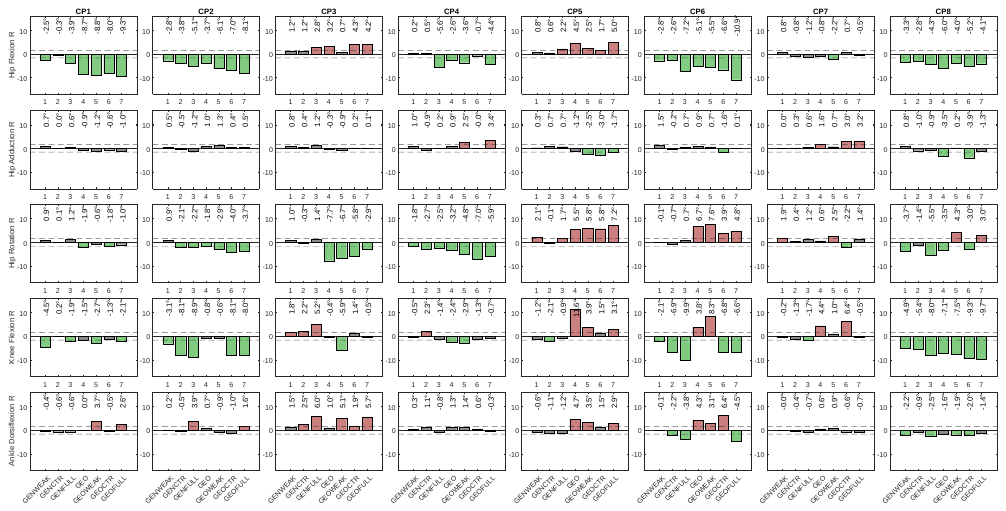

*Fig. S5: Difference in RMSD between simulations and experimental data between personalized models and the reference model GEN for all patients and studied degrees of freedom on the right side. A green bar indicates an assumed contribution of the modeled impairment to altered gait. A red bar indicates modeled impairments are assumed to not contribute to the observed gait pattern. GEN is the generic scaled model, GENWEAK is GEN with weakness, GENCTR is GEN with contractures, GENFULL is GEN with both weakness and contractures. GEO is the model with MRI-based deformities, GEOWEAK is GEO with weakness, GEOCTR is GEO with contractures, and GEOFULL is GEO with both weakness and contractures. GEN represents how a typically developing individual with the same dimensions as the patient would walk.*

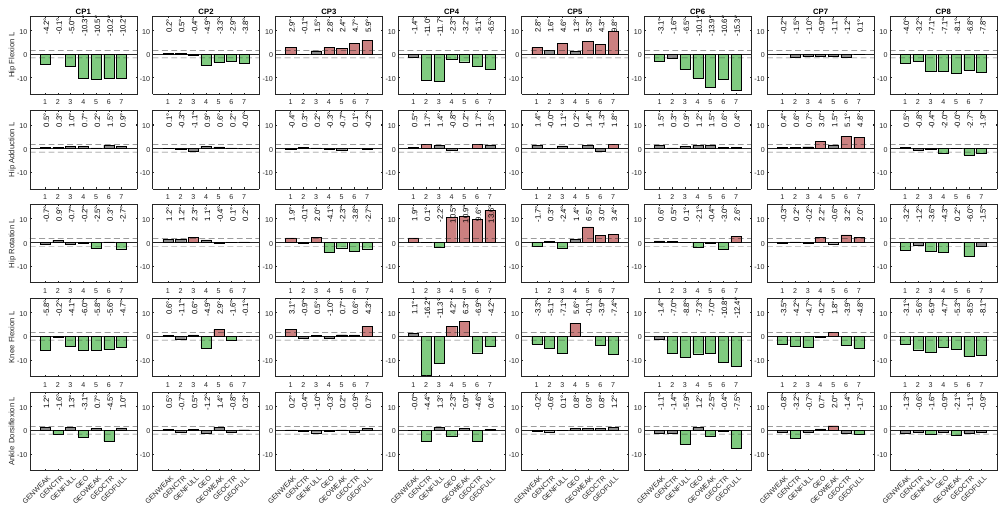

*Fig. S6: Difference in RMSD between simulations and experimental data between personalized models and the reference model GEN for all patients and studied degrees of freedom on the left side. A green bar indicates an assumed contribution of the modeled impairment to altered gait. A red bar indicates modeled impairments are assumed to not contribute to the observed gait pattern. GEN is the generic scaled model, GENWEAK is GEN with weakness, GENCTR is GEN with contractures, GENFULL is GEN with both weakness and contractures. GEO is the model with MRI-based deformities, GEOWEAK is GEO with weakness, GEOCTR is GEO with contractures, and GEOFULL is GEO with both weakness and contractures. GEN represents how a typically developing individual with the same dimensions as the patient would walk.*

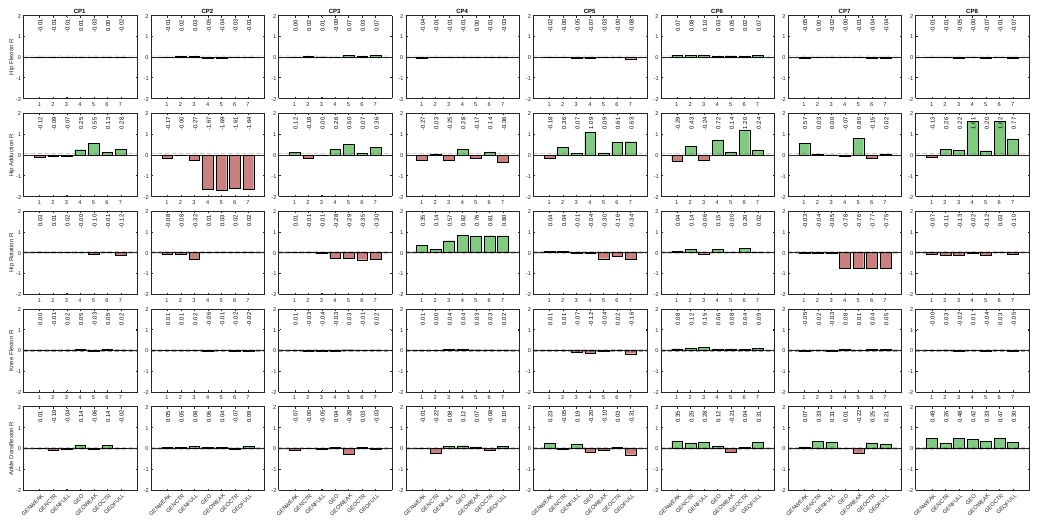

*Fig. S7: Difference in r between personalized models and the reference model GEN for all patients and studied degrees of freedom on the right. A green bar indicates an assumed contribution of the modeled impairment to altered gait. A red bar indicates modeled impairments are assumed to not contribute to the observed gait pattern. GEN is the generic scaled model, GENWEAK is GEN with weakness, GENCTR is GEN with contractures, GENFULL is GEN with both weakness and contractures. GEO is the model with MRI-based deformities, GEOWEAK is GEO with weakness, GEOCTR is GEO with contractures, and GEOFULL is GEO with both weakness and contractures. GEN represents how a typically developing individual with the same dimensions as the patient would walk.*

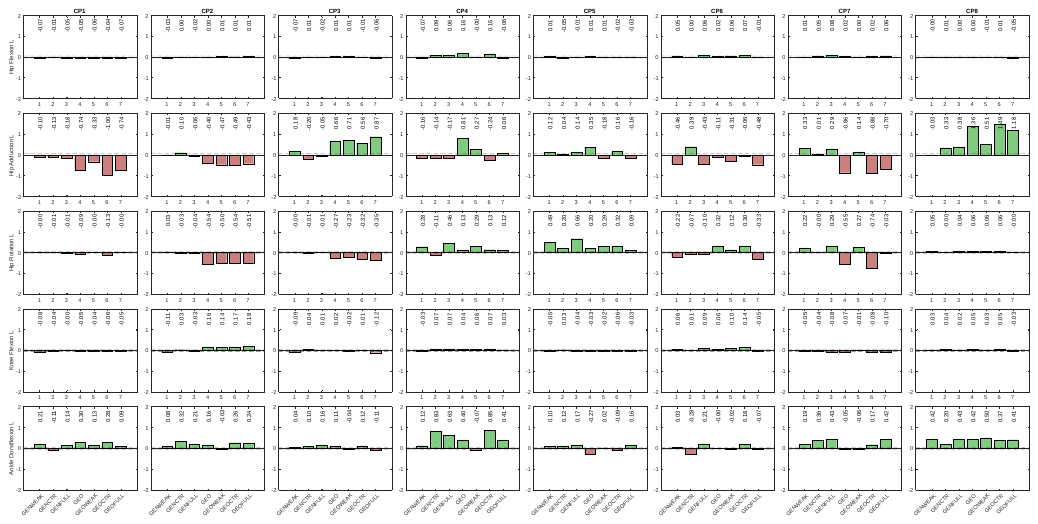

*Fig. S8: Difference in r between personalized models and the reference model GEN for all patients and studied degrees of freedom on the left. A green bar indicates an assumed contribution of the modeled impairment to altered gait. A red bar indicates modeled impairments are assumed to not contribute to the observed gait pattern. GEN is the generic scaled model, GENWEAK is GEN with weakness, GENCTR is GEN with contractures, GENFULL is GEN with both weakness and contractures. GEO is the model with MRI-based deformities, GEOWEAK is GEO with weakness, GEOCTR is GEO with contractures, and GEOFULL is GEO with both weakness and contractures. GEN represents how a typically developing individual with the same dimensions as the patient would walk.*
